# Derivation and analysis of human somatic sensory neuron subtypes facilitated through fluorescent hPSC reporters

**DOI:** 10.1101/2025.02.25.640131

**Authors:** Eti Malka-Gibor, Katherine M. Oliver, Shirley Chen, Alejandro Garcia-Diaz, Damian Williams, Barbara Corneo, Joriene C. de Nooij

**Author notes:** These authors contributed equally (Eti Malka-Gibor, Katherine M. Oliver, Shirley Chen).

## Abstract

Peripheral sensory neuropathy (PSN) is associated with several devastating neurological conditions, yet effective strategies to prevent or alleviate the consequences of PSN are nearly non-existent. A major challenge in the development of better therapeutic interventions is the lack of appropriate human model systems. Human induced pluripotent stem cell (hiPSC)-derived somatosensory neurons present a promising strategy to overcome this issue but remain of limited translational utility, in part due to the low efficiency and lack of sensory subtype selectivity of the existing sensory neuron derivation protocols. To improve upon iPSC-based somatosensory disease models, we here describe the generation and validation of a genetic toolset to fluorescently label all or distinct (nociceptor, low threshold mechanoreceptor, and proprioceptor) somatosensory subtypes. These new resources will be transformative for hPSC-based approaches in PSN disease modeling - a critical step towards translating new findings into clinically relevant therapeutic strategies.

## Introduction

Peripheral sensory neuropathy (PSN) is a neurological condition caused by heritable or idiopathic metabolic disease, exposure to toxic substances (including chemotherapeutic agents), or physical nerve damage (Gomatos and Rehman, 2022; Houlden et al., 2004; De Leon et al, 2023; Feldman et al., 2019). PSN patients suffer from chronic pain, paresthesia, or impaired touch sensation or sensory-motor control – conditions that can severely impact quality of life and often result in disability. With an estimated 20 million people afflicted in the US alone, PSN presents an enormous public health and socioeconomic burden which is estimated to increase with the aging population (Gomatos and Rehman, 2022; Mold et al., 2004). Strategies to prevent or alleviate the consequences of PSN are essentially non-existent. In part, this is due to the heterogeneous nature of the trigeminal- and dorsal root ganglion neurons that provide somatosensory feedback (Usoskin et al., 2015; Li et al, 2016; Nguyen et al., 2017, 2021; Sharma et al, 2020; Bhuiyan et al., 2024). In addition, in both preclinical mouse models and post-mortem human dorsal root ganglia (DRG) tissue, the abundance of individual sensory subtypes is often too scarce for rigorous analyses. Together, the neuronal heterogeneity and the limited experimental material make it difficult to examine the mechanistic basis of disease or to systematically evaluate potential interventions. A second major issue is the difficulty in translating animal studies into effective therapeutics for human subjects (Burma et al., 2017, Mogil, 2019; Hill, 2000; Alsaloum et al., 2020). This problem has been attributed to the differences in gene expression profiles between mouse and human somatosensory subtypes (Han et al., 2015; Ray et al., 2018; Rostock et al., 2018; Nguyen et al, 2021). These observations have emphasized the need for better, and in particular, human-based pre-clinical model systems for sensory neuropathies.

Tissues derived from patient iPSCs have long proven their value in disease modeling and drug development, including for disorders related to peripheral sensory neurons (Fujimori et al., 2018; Zhou et al., 2018; Nickolls et al., 2020; Snavely et al., 2022). Yet, despite isolated successes (Namer et al., 2018, Mcdermott et al., 2019, Sun et al., 2020), the translational utility of *in vitro* derived somatosensory neurons has remained limited (Smulders et al., 2024). The reasons for this are severalfold. First, existing strategies to derive or transdifferentiate sensory neurons from iPSCs or fibroblasts, respectively, often do not yield enough mature sensory neurons to permit the screening of new therapeutic compounds at an industrial level (Chambers et al., 2012, Blanchard et al., 2015; Wainger et al., 2015). Second, although neuropathies often show pathological consequences in select sensory subtypes (Koeppen and Mazurkiewicz, 2013; Finsterer and Scorza, 2022; Gonzales-Duarte et al., 2023), most sensory differentiation protocols result in a mixture of sensory neurons. While this perhaps better represents the normal physiological condition, it also complicates the phenotypic analysis of specific sensory subtypes (Chambers et al., 2012, Blanchard et al, 2015; Alshawaf et al., 2018; Saito-Diaz et al., 2021; Deng et al., 2023). This problem is compounded by the lack of validated immunological reagents that can reliably distinguish between human *in vitro* derived sensory neuron subtypes. Some of these difficulties have in part been addressed by newer differentiation strategies that rely on the transient expression of sets of transcription factors in fibroblasts or iPSCs, often in combination with the addition of specific cytokines, to promote a generic or subtype selective sensory neuron identity (Tsunemoto et al., 2018; Nickolls et al., 2020; Schrenk-Siemens et al., 2022; Hulme et al., 2024). While these “induced sensory neurons” (iSNs) offer some advantages, they present a limitation for many sensory neuron developmental disorders, including neurocristopathies, given that they bypass early developmental stages. Another drawback of such models is the need for doxycycline to induce the transcription factors. Given that doxycycline - even at low doses - can adversely affect cell proliferation and mitochondrial function, the use of this agent may inadvertently modify disease phenotypes (Luger et al., 2018; De Boeck and Verfaillie, 2021).

To address the aforementioned limitations in the existing iPSC-based somatosensory disease models, and to facilitate the optimization of sensory neuron differentiation protocols that best resemble their nascent developmental trajectory, we here describe the generation and validation of a genetic reporter toolset for hESC/iPSC-derived sensory neurons. These reporters include an *AVIL:tdTomato* (tdT) reporter, designed to mark all sensory neurons, and *NTRK1:tdT*, *MAFA:tdT*, and *RUNX3:tdT* reporters, to label nociceptive/thermoceptive, low threshold mechanoreceptive and proprioceptive lineages, respectively. Validation of these reporters demonstrates that expression of the tdT fluorescent marker is restricted to the intended target population. The reporter lines can be used in a variety of applications, ranging from optimization of differentiation protocols, visualization of the neurons in multiplex organoids (innervated skin or bone), to isolating the specific sensory subsets using fluorescent activated cell sorting (FACS) for detailed molecular or biochemical analyses. Each reporter is readily generated for different human ESC or iPSC (hPSC) lines, enabling the rapid generation of new model systems for various types of peripheral sensory neuropathy.

Taken together, our collection of hPSC sensory reporter lines offers a versatile new toolkit to optimize sensory neuron differentiation strategies and to permit systematic phenotypic investigations of select sensory neuron subtypes under pathological conditions. These resources will be valuable in applying hPSC-based approaches in the modeling of a vast number of sensory disorders.

## Results

### Generation and validation of a generic sensory neuron AVIL:tdTomato hESC-reporter line

Somatic sensory neurons reside in DRG and trigeminal ganglia (TG) and comprise a myriad of distinct sensory subtypes – each designed to provide specific feedback regarding changes in the external or internal chemical or physical environment (Usoskin et al., 2015; Li et al, 2016; Nguyen et al., 2017, 2021; Sharma et al, 2020; Bhuiyan et al., 2024). Broadly, these sensory neurons can be divided into three main classes typically referred to as small-diameter nociceptive, itch, or temperature-sensitive neurons; low threshold mechanoreceptors (LTMRs) involved in discriminative touch; and proprioceptors, which mainly innervate muscle and provide feedback on body and limb kinetics (Lallemend and Ernfors, 2012; de Nooij, 2023). At spinal segmental levels, these subsets all derive from the neural crest, a transient progenitor population that delaminates from the dorsal-most aspect of the neural tube to form the nascent DRG (Simoes-Costa and Bronner, 2014). The proximity of the neural tube and DRG means that the neurons in these two tissues are exposed to many of the same external growth factors. This similarity is reflected in *in vitro* protocols that aim to generate dorsal spinal or sensory neurons and which rely on many of the same signaling molecules (Chambers et al., 2012; Saito-Diaz et al., 2021; Duval et al., 2019; Gupta et al., 2022). In addition, early sensory progenitors express several transcription factors also found in developing spinal neuron subsets (e.g., Islet1, Pou4f1, NGN1, NGN2) (Ma et al., 1999; Dykes et al., 2011; Andrews et al., 2017; Andersen et al., 2023). To help distinguish between spinal and sensory lineages during the *in vitro* differentiation of somatic sensory neurons, we developed a human embryonic stem cell (hESC) fluorescent reporter line based on the expression of the *AVIL* gene, which encodes Advillin, an actin regulatory protein of the Gelsolin/Villin family (Marks et al., 1998). Advillin is expressed in most if not all rodent and human DRG neurons, in particular during early developmental stages, but not in spinal cord (Supplemental Figure S1A-C) (Hasegawa et al., 2007; Zurborg et al., 2011; Hunter et al., 2018; Nguyen et al., 2021; Russ et al., 2021; Andersen et al., 2023).

To manipulate the *AVIL* locus in RUES2 hESCs, we used a CRISPR/Cas9-mediated gene editing approach to insert a Cre recombinase transgene (Knott and Doudna, 2018; Garcia-Diaz et al., 2020). To ensure high levels of reporter expression, we coupled *AVIL:Cre* expression to Cre-dependent GAGGS-promotor driven reporter expression using the Cre/loxP system (Brault et al., 2007). In addition, we selected tdTomato (tdT) as our reporter to enable immediate visualization of reporter expression using most standard fluorescent microscopes (i.e. without a need for immunological signal amplification). The *AVIL:Cre* driver was generated by targeting the Cre coding sequence (cloned in frame with a P2A internal ribosomal entry site sequence) to exon 6 of the *AVIL* gene, just before the stop codon (Supplemental Figure S1D) (see methods for details). hESC clones with a correctly inserted Cre sequence (12.5% of clones) were identified by genotyping PCR (Supplemental Figure S1E). One hESC *AVIL:Cre* clone was subsequently used to target a *CAGGS:lxp-STOP:lxp-tdTomato* reporter into the *PPP1R12C* locus (also known as the *AAVS1* safe harbor locus) again using a CRISPR/Cas9 strategy as described previously (DeKelver et al., 2010; Garcia Diaz et al., 2020).

### Validation of the AVIL:tdT reporter

To validate the *AVIL:Cre;AAVS1:tdTomato* reporter (hereafter *AVIL:tdT*) we differentiated the RUES2 hESCs harboring this reporter into sensory neurons using a strategy based on previously described protocols with a few modifications (Chambers et al., 2009, 2012; Maury et al. 2015). To emulate the *in vivo* differentiation trajectory of native sensory neurons, in this protocol hESCs are exposed to various signaling molecules (including the SMAD inhibitors LDN, SB, as well as modulators of the Wnt and Notch signaling pathways) (Figure 1A) (Simoes-Costa and Bronner, 2014; Hari et al., 2002; Lee et al., 2004). During the first differentiation steps, hESCs are plated in low-attachment plates to promote the formation of embryoid bodies (EBs). We noted that for RUES2 cells, the addition of BMP4 at four days of *in vitro* culture (DIV4) was essential to induce the expression of PAX3 (a marker of dorsal neuroepithelium) and neural crest identity (Supplemental Figure S2A-G) (Cimadamore et al., 2011; Duval et al., 2019). Under the influence of the various signaling factors, the RUES2 EBs went through several transcriptional stages, marked by the expression of SOX2 (indicative of a general neural identity), PAX3 (indicating a dorsal neural fate), and AP2α and SOX10 (markers for a neural crest cell identity) (Figure 1A, B, Supplemental Figure S2D) (Cimadamore et al., 2011; Simoes-Costa and Bronner 2014). During these early stages we also detected expression of HOX transcription factors (associated with spinal segmental levels), as well as NEUROGENIN (NGN) 1 and 2, transcription factors known to be expressed early in the somatosensory lineage (Supplemental Figure S2H) (Ma et al., 1999).

**Figure 1.**
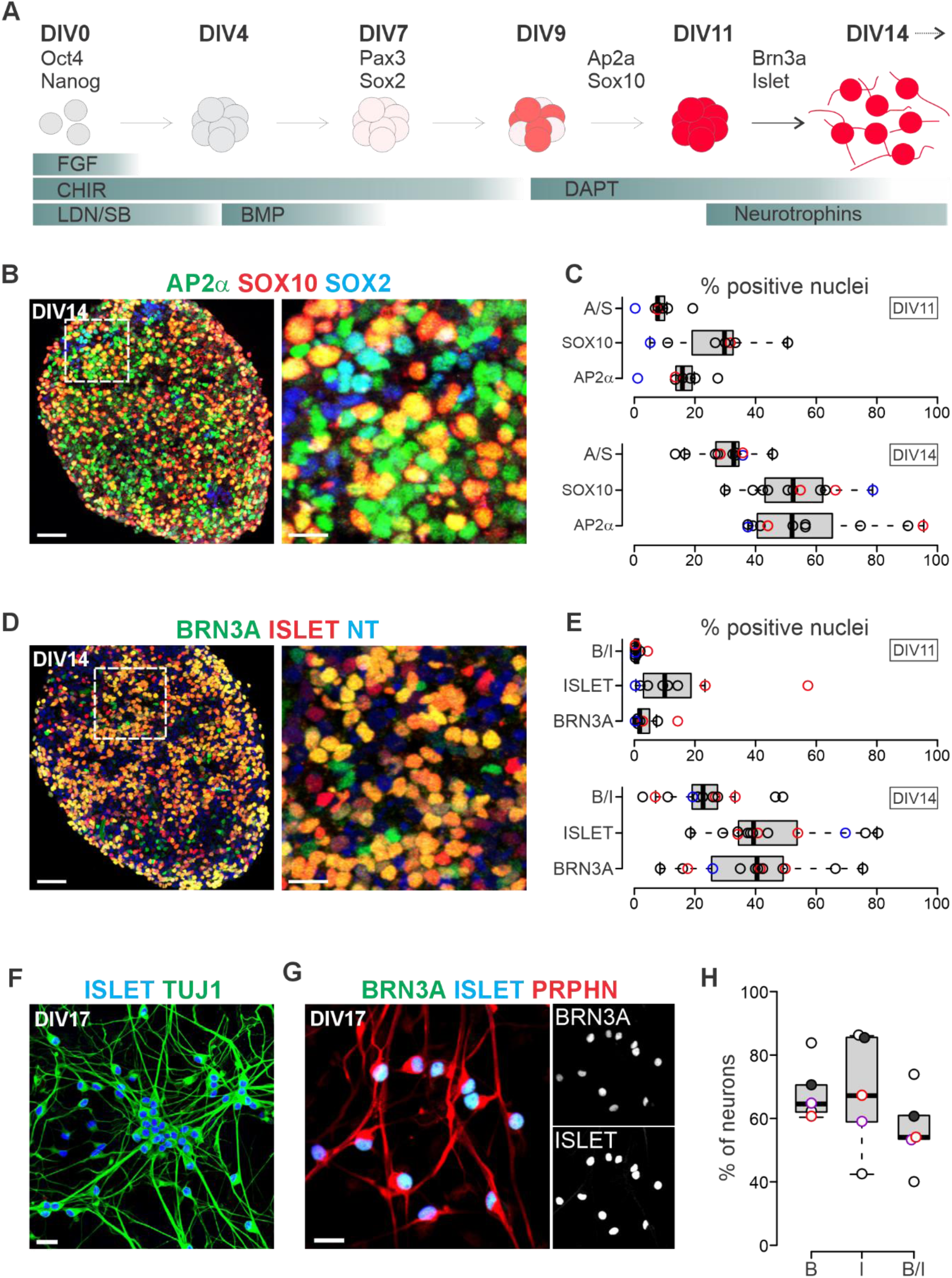
Protocol to derive human sensory neurons from embryonic and induced pluripotent stem cells. A) Schematic diagram describing the main growth factors/cytokines and their time points of application to differentiate human ES cells towards a somatic sensory neuron fate. At DIV0, hESCs are plated in low attachment plates to promote the formation of embryoid bodies (EBs). At DIV14, EBs are dissociated into single cell suspensions and plated in 2D format. Transcription factors that mark specific stages of the differentiation process are indicated. B) Expression of the neural crest markers AP2α and SOX10 and the neural progenitor marker SOX2 in DIV14 EBs. Image on the right is a magnification of the boxed area in the image on the left. C) Percentage of AP2α^+^, SOX10^+^, and AP2α^+^SOX10^+^ cells from the total number of cells in DIV11 and DIV14 EBs. Percentages are estimates based on a machine learning approach. See Methods for details. Data derived from 7 (DIV11) or 11 (DIV14) independent differentiations with at least 2 representative EB images used per data point (differentiation). See below for a legend to the data point color code. D) Expression of the post-mitotic sensory neuron markers BRN3A and ISLET, and the general neuronal marker Neurotrace (NT) in DIV14 EBs. Image on the right is a magnification of the boxed area in the image on the left. (The ISLET antibody used in these experiments recognizes both ISLET 1 and 2.) E) Percentage of BRN3A^+^, ISLET^+^, and BRN3A^+^,ISLET^+^ neurons from the total number of cells in DIV11 and DIV14 EBs. Percentages are estimates based on a machine learning approach using Image J. See Methods for details. Data derived from 8 (DIV11) or 13 (DIV14) independent differentiations with at least 2 representative EB images used per data point. See below for a legend to the data point color code. F) Expression of ISLET and the neuronal marker TUJ1 at DIV17, shortly after plating the dissociated EBs. G) Expression of BRN3A, ISLET, and the peripheral neuronal marker PERIPHERIN at DIV17. H) Percentage of BRN3A^+^, ISLET^+^, and BRN3A^+^,ISLET^+^ post-mitotic sensory neurons from the total number of cells/neurons in DIV17 cultures. Data obtained from 5 independent differentiations; total number of neurons counted/experiment was 166, 203, 556, 268, and 278. Scale: 50 μm (1B, 1D left), 10 μm (1B, 1D right), 20 μm (1F, 1G). Black circles (in 1C, 1E, 1H) represent data obtained from differentiations with CHIR at DIV0-7, 10ng/ml BMP4 at DIV4-7, DAPT at DIV9-14, and GF (NT3, BDNF, GDNF, NGF; 10 ng/ml each) added starting from DIV 11 and onwards. Red circles (in 1C, 1E, 1H) represent data from differentiations with DAPT added on DIV7 (instead of DIV9) and with growth factors (NT3, BDNF, GDNF, NGF) added on DIV9 (instead of DIV11); Blue circles (in 1C, 1E) represent data obtained from differentiations with CHIR prolonged till DIV11 (instead of DIV7). Black solid circles (in 1H) represent data obtained from a differentiation similar to black open circles in 1C, 1E but with growth factors (NT3, BDNF, GDNF, NGF) added on DIV9 (instead of DIV11). Purple circles (in 1H) represent data obtained from differentiations with BMP4 prolonged till DIV11 (instead of DIV7) and DAPT addition delayed till DIV11 (instead of DIV9).

By DIV11-14, EBs start to express the transcription factors POU4F1 (encoding BRN3A) and ISLET1/2 (Figure 1D). (Unless specified otherwise, the ISLET antibody used in our experiments recognized both ISLET1 and ISLET2). Co-expression of BRN3A and ISLET is known to mark a post-mitotic sensory identity (Dykes et al., 2011). To estimate the relative differentiation efficiencies, we developed a FIJI-based machine learning (ML) approach (see methods for details) to quantify the numbers of SOX10, AP2α, BRN3A, and ISLET expressing cells within EB tissue sections. Based on these ML analyses, we estimate that at DIV11, 15.7% (± 3.0% s.e.m.) of EB cells express AP2α, 26.9% (± 5.7%) SOX10, and 8.8% (± 2.2%) co-express both (Figure 1C). Co-expression of BRN3A and ISLET1 is still very low at that stage: 3.9% (± 1.7%) of cells express BRN3A, 15.2% (± 6.6%) ISLET, and just 1.1% (± 0.5%) co-express BRN3A and ISLET (Figure 1E). However, by DIV14, EBs on average have 39.1% (± 5.3%) BRN3A^+^ cells, 45.9% (± 5.2%) ISLET^+^ cells, and 24.1% (± 3.4%) of cells that co-express BRN3A and ISLET (Figure 1D, E). We observed similar levels of AP2α/SOX10 and BRN3A/ISLET expression in four other stem cell lines (hESC or iPSC) that we differentiated using similar protocols (Supplemental Figure S3A, B). Thus, our differentiation protocol is robust when assessing markers for neural crest and post-mitotic sensory neurons.

Upon dissociation of the EBs and replating of the differentiated cells at DIV14, we found that most cells assume a neuronal morphology as they extend neurites shortly after plating. In addition, by DIV17 the majority of BRN3A^+^ and ISLET^+^ neurons co-express class III β-TUBULIN (TUJ1), a neuronal-specific tubulin, and PERIPHERIN, an intermediate filament protein predominantly expressed in peripheral neurons (Figure 1F, G) (Lewis and Cohan, 1988; Holford et al., 1994; Fornaro et al., 2008). At DIV17, the percentage (± s.e.m.) of TUJ1^+^ or PERIPHERIN^+^ neurons positive for BRN3A^+^ neurons is 68.3 ± 4.3%, the percentage of ISLET^+^ neurons is 68.0 ± 8.3%, and the percentage of BRN3A^+^ISLET^+^ neurons is 56.6 ± 5.6% (Figure 1H). Based on these observations, we conclude that our protocol differentiates hESCs into BRN3A^+^ISLET^+^ sensory neurons with relative high efficiency.

Employing this sensory neuron differentiation strategy, we next validated our newly generated *AVIL:tdT* hESC reporter line. Consistent with the expression of ADVILLIN in differentiated mouse and human neurons (Supplemental Figure S1A, B), we found that starting from DIV20-23, *AVIL:tdT* hESC-derived sensory neurons express tdT with the number of tdT^+^ neurons increasing with extended culture (Figure 2A-C). At early stages, very few neurons show tdT expression (0.6% at DIV18). By DIV30, 24.5% (± 2.7%) of neurons express tdT, and by DIV60 this increases to 48.4% (± 5.6%) (Figure 2B, C). We also explored if small modifications to our protocol (e.g., the concentration of BMP4 added, the addition of retinoic acid [RA], the timing and concentration of growth factors [NT3, BDNF, GDNF, or NGF]) would result in higher yields of AVIL:tdT^+^ neurons, but thus far none of these alterations consistently did so (Figure 2C). Using our standard protocol, we find that nearly 100% of tdT^+^ neurons co-express both BRN3A and ISLET, confirming their somatosensory neuron identity (Figure 2D, E). Only extremely rarely did we observe tdT^+^ cells with non-neuronal morphology indicative of ectopic activation of the *AVIL:tdT* reporter (not shown). Interestingly, although all tdT^+^ neurons co-express BRN3A and ISLET, not all BRN3A^+^ISLET^+^ express tdT; at DIV30, we observed tdT in 57.7% (± 5.9%) of BRN3A^+^ISLET^+^ neurons (Figure 2D, F).

**Figure 2.**
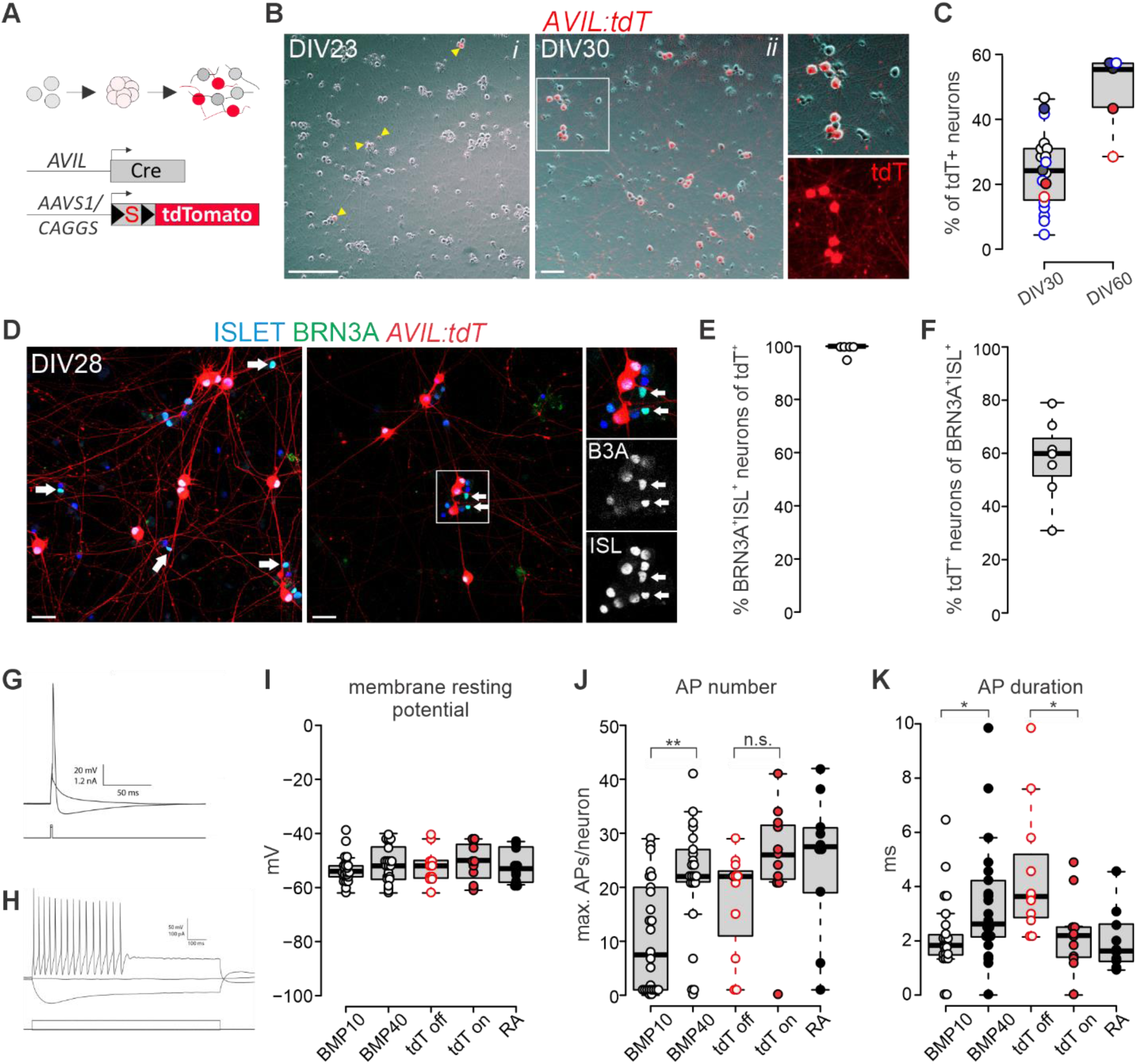
Validation of the *AVIL:tdT* hESC reporter line. A) Genetic strategy to mark developing sensory neurons with the tdTomato (tdT) fluorescent protein. In hESCs that combine an *AVIL:Cre* allele with a Cre-dependent and CAGGs driven tdT reporter targeted to the *AAVS1* locus, emerging sensory neurons that express *ADVILLIN* will be permanently marked by tdT. B) Expression of tdT in *(i)* DIV23 and *(ii)* DIV30 sensory neurons. Boxed area in *(ii)* is enlarged in right most panels to visualize tdT^+^ neurites in addition to sensory neuron cell bodies. C) Percentage of tdT^+^ *AVIL:tdT* hESC neurons at DIV30 and DIV60 across various differentiation protocols (slight modifications of our baseline protocol as described above). DIV30 includes data from DIV28-32; DIV60 includes data from DIV58-60. Data obtained from 19 (DIV30) or 5 (DIV60) independent experiments with the total number of neurons counted per experiment ranging between 48 and 940 (average/experiment for DIV30 is 362.7 ± 51.7 neurons; average for DIV60 is 196.6 ± 35.5 neurons). Black circles represent data from differentiations with CHIR at DIV0-7, BMP4 (10ng/ml) at DIV4-7, DAPT at DIV9-14 and growth factors (NT3, BDNF, GDNF, NGF) 10ng/ml from DIV11-14, and thereafter 25 (NT3), 10 (BDNF), 20 (GDNF) or 50 (NGF) ng/ml till DIV30. Dark gray solid circle represents data from a differentiation similar to open black circles but with 40 ng/ml BMP4 (instead of 10 ng/ml). Solid Black circle represents data from a differentiation with CHIR at DIV0-7, RA (100nM) at DIV2-4, BMP4 (10ng/ml) at DIV4-7, DAPT at DIV7-14, NT3 (25 ng/ml) at DIV9-11, and NGF (50 ng/ml) and GDNF (10 ng/ml) from DIV11-60. Blue circles represent data from differentiations with CHIR at DIV0-4, RA (100nM) at DIV2-4, BMP4 (40ng/ml) at DIV4-7, DAPT at DIV4-14 and growth factors (NT3, BDNF, GDNF, NGF) 10ng/ml from DIV11-14, and thereafter 25 (NT3), 20 (BDNF), 10 (GDNF) or 50 (NGF) ng/ml till DIV30 or DIV60. Solid blue circles represent data from differentiations identical to blue circles but with NT3 and BDNF (both 10 ng/ml) between DIV11-14, GDNF and NGF (both 10 ng/ml) between DIV11-14 and thereafter 25 (NT3), 20 (BDNF), 10 (GDNF) or 50 (NGF) ng/ml till DIV30 or DIV60. Red circles represent data from differentiations similar to blue circles but without RA. Solid Red circles represent data from differentiations identical to solid blue circles but without RA. D) Expression of tdT, BRN3A, and ISLET in *AVIL:tdT^+^* hESC-derived sensory neurons at DIV30. Box indicates enlarged area in right-most panel. Arrows indicate BRN3A^+^, ISLET^+^ neurons that lack tdT expression. E) Percentage of tdT^+^ neurons that co-express BRN3A and ISLET at DIV30. Differentiation conditions identical as described for black open circle data points in (C). F) Percentage of BRN3A^+^ISLET^+^ neurons that co-express tdT at DIV30. Differentiation conditions identical as described for black open circle data points in (C). G) Representative whole cell current clamp recordings showing a subthreshold depolarization evoked by a 2 ms current injection overlaid with an action potential evoked by a higher amplitude current. Current injections at DIV30 elicit action potentials in the vast majority of neurons. H) Overlay of membrane potential traces from the same cell following a 1 s hyperpolarizing or depolarizing current injection. A train of action potentials is evoked by the depolarizing current step. A ‘sag’ in membrane potential is clearly visible when the cell is hyperpolarized. I-K) Electrophysiological charactersitics of hESC-derived sensory neurons treated with 10 or 40 ng/ml BMP4 at DIV4-7. Individual neurons are represented as circles. Neurons derived from EBs treated with 40 ng/ml BMP4 at DIV4-7 are further segregated based on status of tdT expression: tdT^off^ (open red circles) and tdT^+^ (solid red circles). Solid black circles (RA) represent neurons derived from a differentiation protocol with a shortened CHIR (DIV0-4), the inclusion of Retinoic Acid (100nM at DIV2-4), 40 ng/ml of BMP4 (DIV4-7), and earlier DAPT treatment (DIV4-14). Scale: 200 μm (B*i*), 100 μm (B*ii*), 20μm (D).

To determine if the absence of tdT in BRN3A^+^ISLET^+^ neurons is a consequence of an immature sensory neuron identity, we explored the physiological properties of tdT^+^ and tdT^off^ neurons. To do so, we performed whole-cell patch clamp recordings on DIV30 sensory neurons and measured various physiological features that develop with increasing neuronal maturation, including membrane resting potential, membrane resistance, and repetitive firing properties (Figure 2G-K, Supplemental Figure S4A-E).

We observed an average membrane resting potential of -52.4 (± 0.8) mV and found that the vast majority (55/58) of neurons produced an action potential (AP) upon current injection (Figure 2G, I). In addition, most (>75%) of the responding neurons showed repetitive firing with an average of 17.5 (± 1.7) spikes during a 1 s current step at an amplitude which elicited maximum firing (Figure 2H, J). Yet, while we noted a significant difference in repetitive firing and AP duration between neurons that were exposed to higher levels of BMP4 (40 vs 10 ng/ml at DIV4-7), we were unable to detect a significant difference in firing rate, membrane resting potential, membrane resistance, or the presence of an H-current when specifically comparing tdT^+^ and tdT^off^ neurons (all exposed to 40 ng/ml BMP4) (Figure 2I, J, Supplemental Figure S4A, C). However, DIV30 tdT^+^ neurons showed a significantly shorter AP duration (p<0.05, Mann-Whitney Rank Sum Test) (Figure 2K). This suggests that a possible explanation for the presence of tdT^off^, BRN3A^+^ISL^+^ neurons could relate to the maturation state of their membrane or intrinsic firing properties (Lechner and Lewin, 2009; Gao and Ziskind-Conhaim, 1998). We also observed an increase in repetitive firing and shortened AP duration in neurons that were exposed to Retinoic Acid (RA; 100nM at DIV2-4) during the differentiation process (Figure 2J, K). While we have not yet correlated this with increased AVIL:tdT expression, this could indicate that RA may accelerate the neuronal maturation process. Interestingly, when performing these recordings, nearly 60% of all neurons exhibited a small shoulder on the downstroke of the AP (Supplemental Figure S4B, C), a feature more often associated with small diameter nociceptive or temperature-sensitive neurons (Traub and Mendell, 1988; Lechner and Lewin, 2009). Considering that the differentiation protocol we used resembles the protocol by Chambers et al., which yields mostly nociceptor-like neurons (Chambers et al., 2012), we also tested responses to typical nociceptive stimuli, including ATP, capsaicin, and icillin. However, while we observed robust responses to potassium chloride and α,β-methylene-ATP, a P2X3 receptor agonist (Cook et al., 1997; Inoue and Tsuda, 2021), neither *AVIL:tdT^+^* nor *AVIL:tdT^off^* neurons displayed any discernable neural excitation in response to capsaicin or icillin (Supplemental Figure S4D, E, and data not shown).

Lastly, we tested our ability to use the *AVIL:tdT* reporter to purify the neurons for downstream analyses using FACS. We find that dissociated neurons can be isolated with high efficiency and can be replated for axonal regeneration studies (Supplemental Figure S4F, G). Together, these studies demonstrate the utility of the *AVIL:tdT* reporter in optimizing the generation, as well as the phenotypic and electrophysiological analysis of *in vitro* hESC- or iPSC*-*derived somatic sensory neurons.

### Generation and validation of a NTRK1:tdT reporter line

Small diameter nociceptive, thermoceptive, and purinergic sensory neurons together constitute the largest DRG sensory neuron subset and are associated with acute pain, itch, and many chronic pain disorders (Woolf and Ma, 2007). The present need for new, non-opioid analgesics has exacerbated a push for better – and in particular human - model systems to explore and evaluate such new pain medications (Cohen et al., 2021). Despite considerable efforts, to this date, protocols that consistently yield high percentages of mature (capsaicin-, menthol-, or icillin-responsive) nociceptive neurons remain lacking (Chambers et al., 2012. Deng et al., 2023, Smulders et al., 2024). Therefore, to help facilitate new protocol development, we also generated a tdT reporter marking small-diameter nociceptive/temperature-sensitive sensory neurons.

In both mouse and human, developing nociceptive neurons initially express *Ntrk1*, which encodes the Tropomyosin receptor kinase A (TrkA; the receptor for Nerve growth factor [NGF]) (Figure 3A) (Chen et al., 2006; Nguyen et al., 2021; Lu et al., 2024). This suggested that the *NTRK1* locus could offer an opportunity for a genetic reporter insertion in hESCs. In mice, TrkA expression is ultimately extinguished from the non-peptidergic nociceptive population and only maintained in peptidergic nociceptive neurons (Molliver et al., 1997; Chen et al., 2006; Gascon et al., 2010). Therefore, to best reflect *NTRK1* expression dynamics, we coupled tdT expression directly to the *NTRK1* transcript (using a P2A internal ribosomal entry sequence) (Figure 3B). Similar to *AVIL:tdT*, *NTRK1:tdT* hESCs were generated using CRISPR/Cas9 gene editing, and *NTRK1:tdT* clones were identified by genotyping PCR (Figure 3B). Considering that TrkA signaling is essential for nociceptor/thermoceptor survival, we inserted the tdT just before the translational STOP codon such that the TRKA coding sequence remained nearly intact (Figure 3B).

**Figure 3.**
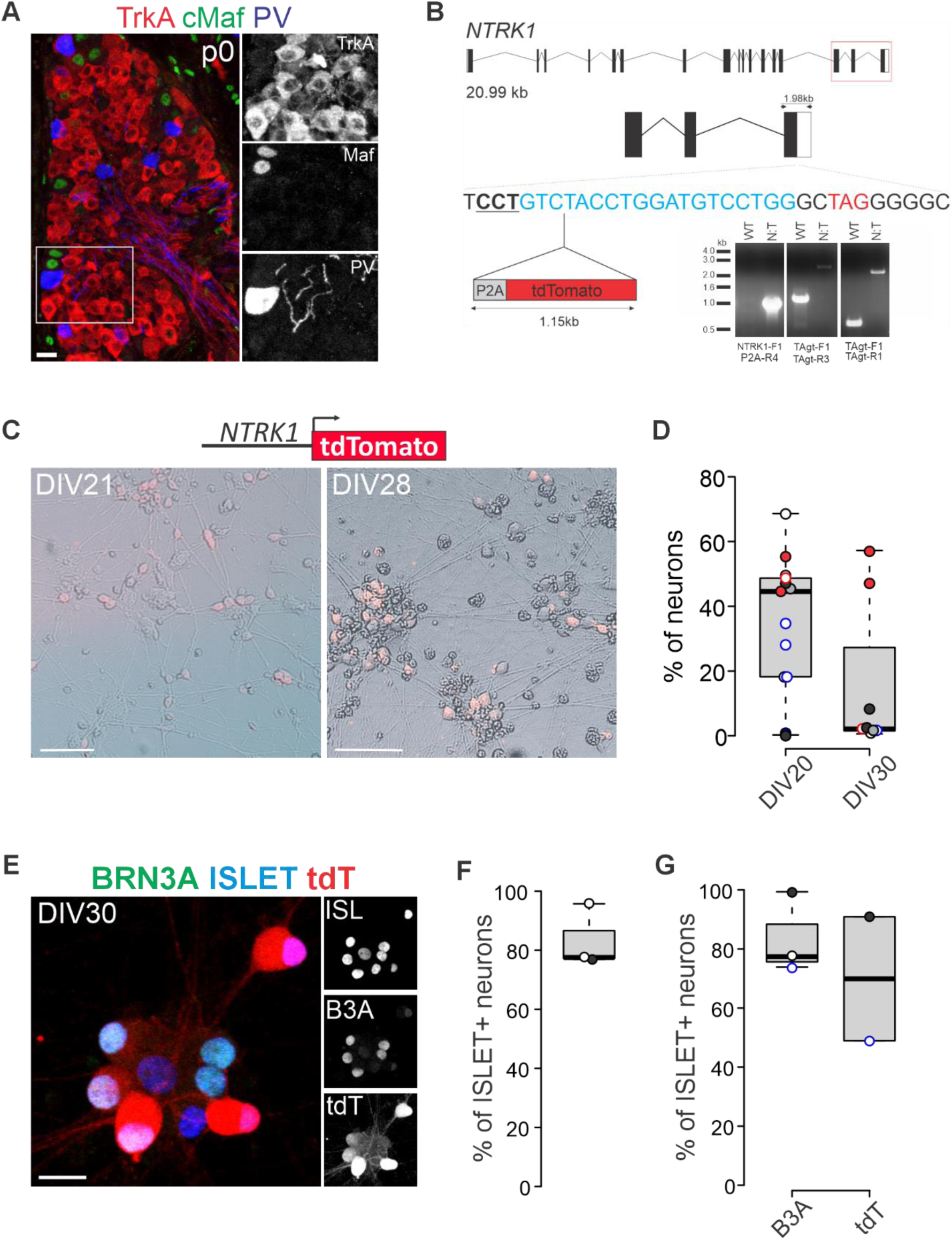
Generation and validation of a *NTRK1:tdT* hESC reporter line. A) Expression of TrkA, cMaf, and Parvalbumin (PV) in P0 mouse DRG. Box in left panel shows area enlarged in panels on the right. B) Strategies to generate the *NTRK1:tdT* reporter using CRISPR/Cas9 gene editing and to genotype a successful gene-edited reporter. See methods for details. C) NTRK1:tdT^+^ sensory neurons in DIV21 and DIV28 cultures. D) Percentage of tdT^+^ NTRK1:tdT hESC neurons at DIV20 and DIV30 across various differentiation protocols. DIV20 includes data from DIV17-22; DIV30 includes data from DIV28-30. Data obtained using three different *NTRK1:tdT* hESC clones (F5, G2, H7) from 13 (DIV20) or 8 (DIV30) independent experiments, with the total number of neurons counted per experiment ranging between 78 and 2035 (average/experiment for DIV20 is 696.8 ± 155.7 neurons; average for DIV30 is 483.1 ± 156.7 neurons). Black open circles represent data from differentiations with CHIR at DIV0-11, RA (100nM) at DIV2-4, BMP4 (40ng/ml) at DIV4-7, DAPT at DIV11-14 and growth factors (GDNF, NGF) each 10ng/ml from DIV11 onwards. Dark gray solid circle represent data from a differentiation similar to solid black circle but with CHIR from DIV0-7, and DAPT from DIV7-14. Solid Black circle represents data from differentiations identical to gray solid circle but with the addition of NT3 (25 ng/ml) at DIV9-11, and NGF increased to 50 ng/ml at DIV11 and onwards. Blue open circles represent data from differentiations with CHIR at DIV0-7, no RA, BMP4 (40ng/ml) at DIV4-7, DAPT at DIV9-14, and GFs (NT3, BDNF, GDNF, NGF) each at 10 ng/ml starting from DIV11. Blue solid circles represent data from differentiations with CHIR at DIV0-11, RA (100nM) at DIV2-4, BMP4 (40ng/ml) at DIV4-7, DAPT at DIV9-14, and GFs at 25 (NT3), 20 (BDNF), 10 (GDNF), or 50 (NGF) ng/ml starting from DIV11. Red open circles represent data from differentiations with CHIR at DIV0-7, RA (100nM) at DIV2-4, BMP4 (40ng/ml) at DIV4-7, DAPT at DIV9-14, and GFs at 10 (GDNF) and 50 (NGF) ng/ml starting from DIV11. Red solid circles represent data from differentiations identical to red open circles but with DAPT from DIV7-16. E) NTRK1:tdT^+^ neurons co-express ISLET and BRN3A F) Percentage of ISLET^+^DAPI^+^ neurons in DIV30 NTRK1:tdT^+^ sensory neurons. DIV30 includes data from DIV30 and 31. Data obtained using one *NTRK1:tdT* hESC clone (F5) from 3 independent experiments, and with the total number of neurons counted per experiment ranging between 176 and 472 neurons. Black open circles represent data from differentiations with CHIR at DIV0-7, RA (100nM) at DIV2-4, BMP4 (40ng/ml) at DIV4-7, DAPT at DIV9-14 and GFs at 25 (NT3), 20 (BDNF), 10 (GDNF), and 50 (NGF) ng/ml starting from DIV11. Black solid circle represents data from a differentiation identical to black open circles but with the DAPT from DIV11-14, GF at 25 (NT3), 20 (BDNF), 10 (GDNF), and 50 (NGF) ng/ml between DIV9-14, followed by only GDNF and NGF thereafter (in same concentrations as DIV9-14). G) Percentage of ISLET^+^ neurons that co-express BRN3A or tdT in DIV30 NTRK1:tdT hESC-derived sensory neurons. DIV30 includes data from DIV30 and 31. Data obtained using one NTRK1:tdT hESC clone (F5) from three (BRN3A) or two (tdT) independent experiments, and with the total number of neurons counted per experiment ranging between 97 and 209 (BRN3A) or 88 and 209 (tdT). Black open circles, and black solid circles represent data from differentiations as described for (F). Blue open circles represent data from a differentiation with CHIR at DIV0-7, no RA, BMP4 (40ng/ml) at DIV4-7, DAPT at DIV9-14, GFs at 25 (NT3), 20 (BDNF), 10 (GDNF), and 50 (NGF) ng/ml at DIV11-14, and just NGF (50 ng/ml) and GDNF (10 ng/ml) starting at DIV14 and onwards. Scale: 20 μm (A, E), 50 μm (C-left), 100 μm (C-right).

Upon differentiation of *NTRK1:tdT* hESCs into sensory neurons (using the same general protocol as described for *AVIL:tdT* neurons) (Figure 1A), we detected tdT fluorescence as early as DIV17 and the average percentage of tdT^+^ neurons was 35.4% ± 5.8 % by DIV20 (Figure 3C, D). For one of our *NTRK1:tdT* hESCs lines, the percentage of tdT^+^ neurons subsequently declined to just 2.6% ± 1.1 % of neurons by DIV30, while in other lines the percentage remained the same or increased (Figure 3C, D). Loss of *TRKA* expression has similarly been observed in other sensory neuron differentiation protocols (Chambers et al., 2012; Deng et al., 2023) and possibly reflects a switch to a non-peptidergic nociceptive/thermoceptive phenotype (Molliver et al., 1997; Chen et al., 2006; Gascon et al., 2010) although this remains to be tested. Alternatively, given that some of our lines did maintain TRKA expression at later stages, it is also possible that intrinsic differences between hESC lines may influence the ability of some hESC clones to continue to develop into mature TRKA^+^ neurons. Expression of tdT was nearly always restricted to ISL1^+^ or POU4F1^+^ISLET^+^ neurons (Figure 3E, F). The number of NTRK1:tdT^+^ neurons increased when detecting tdT using immunological staining (Figure 3G), suggesting that at later stages many neurons may still express tdT but at lower levels. Consistently, many neurons from a *NTRK1:tdT* hESC line that showed little to no native tdT fluorescence at this stage, were tdT^+^ after immunolabeling. Together, these data indicate that the *NTRK1:tdT* reporter reliably marks the presence of TRKA^+^ somatosensory neurons. Considering the relative inefficiency of current nociceptor differentiation strategies with respect to their mature physiological properties, more work is required to optimize these protocols. The *NTRK1:tdT* reporter we describe here should help to facilitate this process.

### Generation and validation of mechanoreceptor and proprioceptor reporter lines

Efforts to develop or test new therapeutics that control or reduce neuropathic pain vastly outnumber studies of neuropathies that also, or primarily, involve large caliber sensory neurons. Many of these disorders, such as hereditary sensory and autonomic neuropathy type III (HSAN III), multiple sclerosis (MS), diabetic neuropathy, chemotherapy induced peripheral neuropathy (CIPN) or Friedreich ataxia (FA) (Gomatos et al., 2024; Koeppen and Mazurkiewicz, 2013; Macefield et al., 2010), have similarly devastating consequences for quality of life. An added complication in modeling these large fiber neuropathies is the sparsity of these sensory subtypes in DRG. On average, just 5-8% of MafA^+^ (“mechanoreceptor”) or Parvalbumin (PV)^+^ (“proprioceptor”) neurons are found in DRG, limiting their availability for either mouse or human DRG studies (Figure 3A) (Bourane et al., 2009; Lecoin et al., 2010; Wu et al., 2019; Nguyen et al., 2021). Existing protocols either focus on generating neurons with elevated levels of NTRK3 expression, or rely on doxycycline-inducible expression of sensory neuron or proprioceptor-selective transcription factors (Schrenk-Siemens et al., 2015; Nickolls et al., 2020; Dionisi et al., 2020; Hulme et al., 2024). While these protocols are useful in certain contexts, they lack in specificity and/or in the ability to model developmental disorders. Therefore, protocols to derive these neuronal subtypes for disease modeling are as much in need as nociceptor-differentiation protocols. To facilitate the development of hESC/iPSC differentiation protocols that bias sensory neurons towards a mechanoreceptor or proprioceptor phenotype, we also developed *MAFA:tdT* and *RUNX3:tdT* reporter lines, respectively.

MafA, a basic leucine-zipper (bZIP) transcription factor of the AP1 family, marks multiple classes of low threshold non-nociceptive mechanoreceptive sensory neurons in mouse and human, including Meissner afferents, Merkel cell afferents and longitudinal lanceolate endings (Bourane et al., 2009; Lecoin et al., 2010; Wende et al., 2012). MafA is not expressed in sympathetic neurons but labels a small subset of spinal interneurons (Bikoff et al., 2016). Runx3, in turn, is a Runt-domain transcription factor, which is nearly exclusively expressed in proprioceptive muscle afferents that supply muscle spindles and Golgi tendon organs (Inoue et al., 2002; Levanon et al., 2002; Kramer et al. 2006). Runx3 expression persists in adult proprioceptive neurons and is not expressed in spinal or sympathetic neurons. Based on these prior observations, we hypothesized that *MAFA:tdT* and *RUNX3:tdT* reporters should similarly delineate the majority of human low threshold mechanoreceptors and proprioceptors, respectively.

Generation of *MAFA:tdT* and *RUNX3:tdT* reporters followed the same general strategy as described for *NTRK1:tdT*, by coupling tdT expression to the endogenous *MAFA* or *RUNX3* transcript using a P2A IRES sequence and with minimal disruption of the *MAFA* or *RUNX3* coding sequences (Supplemental Figure S5A, B). To validate the *MAFA:tdT* and *RUNX3:tdT* reporter lines, we explored several adaptations to our “nociceptor” protocol in an attempt to bias neurons towards a mechanoreceptive/proprioceptive fate. Previous studies had noted that reduced WNT signaling coupled to early DAPT exposure increased expression of NTRK2 (Chambers et al., 2012), a marker typically associated with low threshold mechanoreceptors (Kramer et al., 2006; Perez-Pinera et al., 2008). This led us to explore these modifications in our own protocol. When implementing these changes, we obtained similar efficiencies with respect to POU4F1^+^ISLET^+^ sensory neurons in DIV11 EBs when compared to our previously described protocol (Figure 4A, B, Supplemental Figure S5C-F). Moreover, after dissociating the EBs and replating the neuronal cultures, we detected tdT-expressing neurons for both reporter lines, albeit with vastly different efficiencies (Figure 4C, D, F, G). MAFA:tdT^+^ neurons were readily observed by DIV18 and continued to increase to DIV30 (Figure 4C, D, Supplemental Figure S5H). While we did not have a working MAFA antibody, all MAFA:tdT^+^ neurons co-expressed the close family member cMAF. cMAF is also expressed in mechanoreceptive sensory neurons, therefore indirectly validating the low threshold mechanoreceptive identity of the MAFA:tdT^+^ neurons (Figure 4E, I) (Wende et al., 2012). Consistent with this, nearly all MAFA:tdT^+^ neurons co-expressed TRKB (Figure 4I, J). Though expression of tdT is more sparse in the *RUNX3:tdT* reporter line (but see Supplemental Figure S5G, H), all tdT^+^ neurons co-expressed Runx3 (Figure 4F-H, Supplemental Figure S5G).

**Figure 4.**
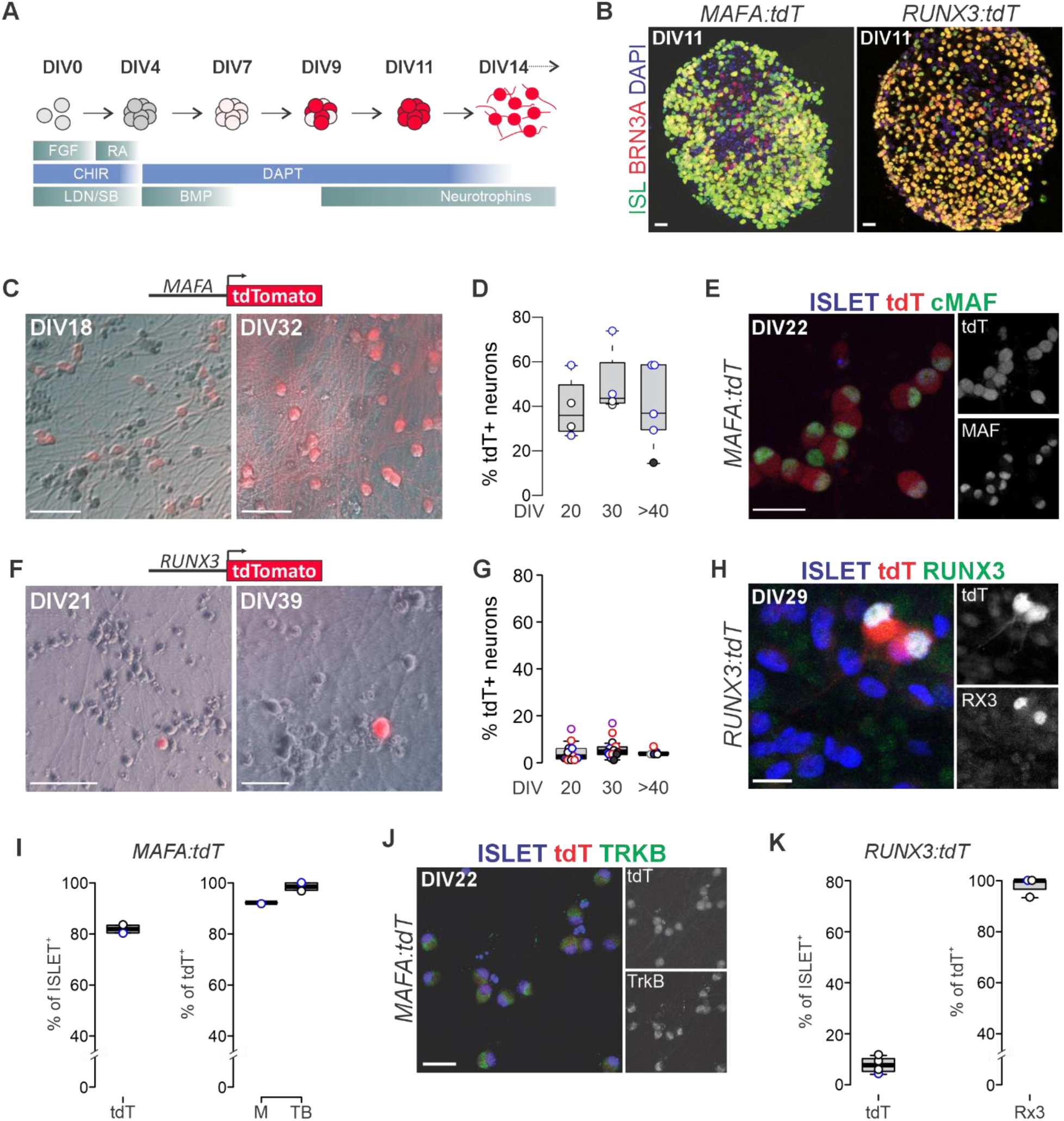
Generation and validation of hESC reporter lines for low threshold mechanoreceptors and proprioceptors. A) Schematic of ESC differentiation protocol to bias neurons toward a mechanoreceptor and/or proprioceptor sensory identity. Protocols diverge only slightly for the two subtypes with the addition of BDNF/GDNF and NT3 neurotrophic factors for mechanoreceptors and proprioceptors, respectively. See Methods for details. B) Expression of BRN3A and ISLET in DIV11 EBs derived from *MAFA:tdT* (mechanoreceptor) and *RUNX3:tdT* (proprioceptor) hESCs. C) Expression of tdT in DIV18 and DIV30 “mechanoreceptor” sensory neurons derived from *MAFA:tdT* hESCs. D) Percentage of tdT^+^ neurons in DIV20, DIV30, and DIV≥40 neural cultures derived from MAFA:tdT hESCs directed towards a mechanoreceptor identity through various differentiation protocols. DIV20 includes data from DIV18-22; DIV30 includes data from DIV28-30; DIV≥40 includes data from DIV37-50. Data obtained using two different MAFA:tdT hESC clones (F10, G10) from 4 (DIV20), 4 (DIV30), or 5 (DIV≥40) independent experiments, and with the total number of neurons counted per experiment ranging between 42 and 360 (average/experiment for DIV20 is 131.5 ± 3.5 neurons; average for DIV30 is 52.5 ± 10.2 neurons; average for DIV≥40 is 161.8 ± 50.4 neurons). Black open circles represent data from differentiations with CHIR at DIV0-4, RA (100nM) at DIV2-4, BMP4 (40ng/ml) at DIV4-7, DAPT at DIV4-14 and GF at 25 (NT3), 20 (BDNF), 10 (GDNF), and 50 (NGF) ng/ml from DIV11 onwards. Black solid circle represents data from a differentiation identical to black open circles but with DAPT from DIV4-11, NT3 (25 ng/ml) between DIV9-11, and only BDNF (20 ng/ml) and GDNF (10 ng/ml) from DIV11 onwards. Blue open circles represent data from differentiations identical to black open circles but without the addition of RA. E) MafA:tdT^+^ neurons co-express ISLET and cMAF at DIV30. F) Expression of tdT in DIV21 and DIV39 “proprioceptor” sensory neurons derived from *RUNX3:tdT* hESCs. G) Percentage of tdT^+^ neurons of the total number of neurons in DIV20, DIV30, and DIV≥40 neurons derived from *Runx3:tdT* hESCs directed towards a proprioceptor identity through various differentiation protocols. DIV20 includes data from DIV17-22; DIV30 includes data from 28-31; DIV≥40 includes data from DIV37-45. Data obtained using three different *RUNX3:tdT* hESC clones (A2, C10, H6) from 11 (DIV20), 13 (DIV30), or 5 (DIV≥40) independent experiments, and with the total number of neurons counted per experiment ranging between 21 and 1009 (average/experiment for DIV20 is 257.5 ± 107.9 neurons; average for DIV30 is 270.0 ± 82.4 neurons; average for DIV≥40 is 345.8 ± 70.6 neurons). Black open circles represent data from differentiations with CHIR at DIV0-4, RA (100nM) at DIV2-4, BMP4 (40ng/ml) at DIV4-7, DAPT at DIV4-14 and growth factors (NT3 at 25 ng/ml; BDNF at 20 ng/ml) from DIV7 onwards. Black solid circles represent data from differentiations identical to black open circles but with DAPT from DIV4-7 and without the addition of BDNF. Gray solid circle represents data from a differentiation identical to black solid circles but with RA extended from DIV2-9. Red open circles are differentiations identical to black open circles but with the growth factors added starting on DIV11 and including GDNF (10 ng/ml) and NGF (50 ng/ml). Red solid circle represents a differentiation identical to red open circles but with the addition of RA (1 mM) between DIV9-11. Blue open circles represent data from differentiations identical to red open circles but without the addition of RA at DIV2-4 (or DIV9-11). Blue solid circle represents data from a differentiation identical to black open circles but without the addition of RA at DIV2-4 (or DIV9-11). Purple open circles represent data from differentiations identical to black solid circles but with DAPT from DIV4-11 and NT3 added at DIV9 and onwards at 25ng/ml) H) RUNX3:tdT^+^ neurons co-express ISLET and RUNX3 at DIV30. I) Percentages of tdT^+^ISLET^+^ (left graph), and MAFA^+^tdT^+^ and TrkB^+^tdT^+^ (right graph) neurons at DIV22. Data obtained using one *MAFA:tdT* clone (F10) from two independent experiments, and with the total number of neurons counted per experiment ranging between 90 and 271 (ISLET) and 67-213 (tdT). Colored data points represent differentiations as in (D). J) MAFA:tdT^+^ mechanoreceptive sensory neurons co-express ISLET and TRKB K) Percentages of tdT^+^ISLET^+^ (left graph), and RUNX3^+^tdT^+^ (right graph) neurons at DIV30. DIV30 includes data from DIV29-30. Data obtained using one *RUNX3:tdT* hESC clones (C10) from 4 (tdT) or 3 (RX3) independent experiments, and with the total number of neurons counted per experiment ranging between 48 and 329 (average/experiment for tdT is 183 ± 61.3 neurons; average for RX3 is 228.0 ± 58.9 neurons). Colored data points represent differentiations as in (G). Scale: 20 μm (B, E, H, J), 100 μm (C, F).

These data confirm that our *MAFA:tdT* and *RUNX3:tdT* reporters recapitulate the endogenous *MAFA* and *RUNX3* expression patterns. Future work is needed to further increase the yields of these *in vitro* generated neuronal subsets, in particular for RUNX3 proprioceptors, and to compare their molecular and physiological properties with those of their nascent *in vivo* counterparts. The *MAFA:tdT* and *RUNX3:tdT* reporters described here should help advance these studies.

## Discussion

Somatic sensory neurons comprise many subtypes that are broadly grouped into high threshold nociceptive-, temperature- or itch-sensitive neurons, low threshold touch mechanoreceptors, and proprioceptors, and are each marked by select molecular determinants that underlie their distinct functions. Despite the urgent need for better preclinical sensory neuropathy models, robust protocols that reliably generate large quantities of these human sensory neuron subtypes largely remain lacking. We here describe the generation and validation of four genetic tools designed to help offset these inherent difficulties in the *in vitro* derivation of somatic sensory neurons from human ESCs and iPSCs. These resources should dramatically facilitate the optimization of current sensory neuron differentiation protocols and will aid in the phenotypic analyses of the individual subtypes in sensory neuropathic disease models.

### Optimizing sensory neuron derivation protocols for disease modeling

Protocols for the derivation of somatic sensory neurons have been described previously (Chambers et al., 2012; Maury et al., 2015, Blanchard et al. 2015; Waigner et al., 2015; Deng et al., 2023; Hulme et al., 2024). Generally, they are based on chemical strategies that involve the successive addition of small molecules or growth factors to mimic the *in vivo* environment of developing sensory neurons, sometimes in combination with the directed expression of sensory neuron transcription factors. However, many of these protocols yield mixed populations of sensory neurons, rendering individual subtypes difficult to isolate for analysis. These limitations present a challenge in disease modeling, given that sensory neuropathies often affect specific sensory neuron subtypes. The sensory neuron reporter lines that we describe here can mitigate these issues at multiple levels. First, hPSC sensory neuron differentiation protocols can be optimized in real-time based on the direct readout of the tdTomato fluorescent reporter. Small molecules, neurotrophic factors, and ECM substrates can all be applied in multiple combinations and their effectiveness assessed through the number of tdT^+^ neurons obtained. In similar manners, myriad other signaling factors can be tested to optimize differentiation protocols for specific classes of TRKA^+^ somatosensory neuron subtypes that project to bone, gut, muscle, or visceral organs instead of skin. Likewise, low threshold mechanoreceptors and proprioceptors may be further directed into distinct subtypes with repeated testing of candidate signaling molecules (Handler et al., 2023; de Nooij and Zampieri, 2023).

Another common issue with many hPSC differentiation protocols is stem cell line specific features. Small intrinsic differences in the properties of ESCs or patient-derived iPSCs (e.g., endogenous BMP4 or Wnt expression levels) may diminish the effectiveness of a given differentiation protocol. As such, genuine improvement in sensory neuron protocol development means a robust protocol that works with at least 50% efficiency for all iPSC/ES lines. The tdT reporters we describe can be readily generated for any other ESC/iPSC line through a simple nucleofection with the donor, guide RNA, and Cas9 expression plasmid (all available upon request). Such an approach permits assessing differentiation efficiencies across multiple ESC/iPSC lines. Once a robust protocol is achieved, the reporter itself may no longer be needed, except for specific applications. Strategies that promote a more “natural” differentiation of sensory neuron subtypes would also be desirable over the use of methods that rely on the doxycycline-mediated inducible expression of transcription factors, given that patient-derived iPSCs in particular could be negatively influenced by the exposure to doxycycline.

A third advantage of the hESC/iPSC reporter lines we developed is that even if a differentiation protocol initially yields a limited number of sensory neurons of the desired subtype, one can isolate these for analyses using FACS, or can focus histological (or other) analyses on the tdT^+^ cell bodies/neurites in the mixed culture. Indeed, the tdT reporters used in these studies are sufficiently bright to permit the use of automated plate-reader based analyses in axonal regeneration studies, toxicity analyses, or neuronal activity measurements using calcium imaging. A related application of the sensory neuron tdT reporter lines is their use in multiplex 3D co-cultures such as innervated skin or bone. Expression of the tdT reporter will enable the direct visualization of sensory axons in such systems.

### Limitations of the individual sensory neuron tdT reporter lines

The reporters we describe here do not mark a single sensory subtype but rather represent broader populations of presumptive TRKA^+^ nociceptors, thermoreceptors, or puriceptors, MAFA^+^ Pacinian-, Meissner-, or Lanceolate ending mechanoreceptors, or RUNX3^+^ groups Ia, Ib, or II proprioceptors or Merkel cell afferents (de Nooij et al., 2013). In addition, TrkA is also expressed in sympathetic neurons, and as such this reporter will require additional validation to ascertain that the tdT^+^ neurons constitute somatic DRG sensory neurons. Therefore, these reporters cannot by themselves be used to definitively mark a select sensory subtype but require secondary validation through alternative strategies.

The *Avil:Cre;CAGGlxp-stop-lxp:tdT* (*AVIL:tdT*) hESC line we generated appears most useful for mature neurons given its gradual increase in expression with prolonged culture. Its Cre/LoxP-dependent tdT reporter offers both advantages and disadvantages. For instance, the CAGGS promoter drives very high levels of tdT expression, further facilitating the visualization of neuronal cell bodies and axons. Another advantage is the versatility of the Cre/LoxP system given that *AVIL:Cre* can easily be coupled to other reporters such as a GCamp calcium indicator, or any specific experimental modulators (e.g. siRNA) to test new hypotheses. A potential disadvantage of the *AVIL:tdT* line is that tdT accumulation could influence neuronal health, an important consideration when used in disease modeling. In addition, the nature of this Cre/LoxP system is such that it permanently labels a neuron even if *ADVILLIN* expression were to be transient and not maintained.

### Applications for NTRK1, MAFA, and RUNX3 reporters beyond somatic sensory neurons

Considering that NTRK1, MAFA and RUNX3 serve important roles in other developmental processes and tissues, the reporters we developed can also be utilized for the *in vitro* modeling of a range of other cell types. As described above, sympathetic neurons similarly rely on NGF for their survival and development and express NTRK1 (Smeyne et al., 1994, Hickman et al., 2018). Likewise, MAFA represents a critical molecular marker for pancreatic beta cells and in which it is required to drive insulin expression (Zhang et al., 2015, Aramata et al., 2007). Lastly, while RUNX3 is largely selective for proprioceptive sensory neurons, outside of the nervous system its expression is required in CD4 and CD8 T-cell lineage selection, is found in hair cell follicles, and has a role in spermatogenesis (Egawa et al., 2007, Raveh et al., 2005, Rahmawati et al., 2023). Clearly, the current set of sensory neuron fluorescent reporter lines will not just be relevant for the somatosensory field, but also serve as a valuable resource for *in vitro* hPSC differentiation and/or disease modeling of various other biological tissues.

## Methods

### hES/iPSC culture and maintenance

Human embryonic stem cell (hESC) and induced pluripotent stem cell (iPSC) lines used for this study include RUES2 (NIHhESC-09-0013; Rockefeller University), H9 (NIHhESC-09-0022; Harvard University), FA0000011 (Patel et al., 2020), FRDA4676 and FRDA68 (both corrected FRDA lines from the Friedreich’s Ataxia Cell Line Repository; UT Southwestern) (Li et al., 2016; Misiorek et al., 2020). Experiments conducted with these ESC and iPSC lines were approved by Columbia University’s Human Embryo and Embryonic Stem Cell Research Committee and the Institutional Review Board from Columbia’s Human Research Protection Office (protocol #AAAT9363). hESCs/iPSCs were cultured on mouse embryonic fibroblast (MEF) feeders [Gibco, #A34180] or feeder-free on Matrigel [Corning BD, #356230] or Cultrex [Biotechne, #BME001-05] coated plates. (Matrigel and Cultrex were prepared as per manufacturer’s directions and stocks were diluted 1:100 with DMEM [Life Technologies, #11960077] to the (1x) working concentration; plates typically were incubated for 45-60 min at 37°C prior to use). Cells on feeders were fed daily with hESC media (DMEM-F12 [Life Technologies, #11320033], 20% Knock-out Serum Replacement (KSR) [Life Technologies, #10828-028], 1X Non Essential Amino Acids (NEAA) [Life Technologies, #11140-050], 1X Penicillin/Streptomycin (P/S) [Life Technologies, #15140122], 1X Glutamine [Life Technologies, #25030081], 0.1 mM 2-Mercaptoethanol [Life Technologies, #21985023] and 4 ng/ml FGF2 [R&D Systems, #233-FB]). hESC media was filtered (0.22 μm, low protein binding filters [Corning, #430767]) prior to adding FGF. Feeder-free cultures were fed with mTeSR™ Plus [STEMCELL Technologies, #100-0274] or mTeSR™1 [STEMCELL Technologies, #85850], with media changes every other day. hESC and mTeSR™1 feeding media were supplemented with additional FGF2 (5 ng/ml) StemBeads [Stemcultures SB-500] for o/w culture (two days without media exchange). When using mTeSR™ Plus, o/w cultures were provided with 2x the normal feeding volume. Cells were passaged using 0.5 mM EDTA [Life Technologies, #15575-020] and mechanical trituration. Rock inhibitor (10 μM) (Y-27632 dihydrochloride) [Tocris, #1254/50], 1x CloneR™2 [STEMCELL Technologies, #100-0691], or 4 μl/ml CEPT (50 nM Chroman 1 [MCE, #1273579-40], 5 μM Emricasan [Selleck Chemicals, #7775], Polyamine supplement (1:1,000) [Tocris, #7739], 0.7 μM Trans-ISRIB [Tocris, #5284]) was added after passaging the cells to increase survival (Chen et al., 2021). All hPSCs were tested routinely for Mycoplasma using the e-Myco™ plus Mycoplasma detection Kit (Intron, #25234). All cells were kept in a normoxic incubator, at 37°C, 5% CO2.

### hESC/iPSC differentiation

hESCs/iPSCs were differentiated into sensory neuron based on a protocol adapted from Chambers et al., 2012, and Maury et al., 2015. hESCs/iPSCs used in differentiations were passaged onto Matrigel or Cultrex-coated feeder-free plates and expanded until 70-80% confluent. Cells were rinsed once and then incubated in 0.5 mM EDTA. After 3 minutes of incubation (at RT), EDTA was aspirated and cells were mechanically triturated in 1 ml of N2B27 media (50% Advanced DMEM/F12 [Life Technologies, #12634028], 50% Neurobasal [Life Technologies, #21103049], 1X P/S [Life Technologies, #15140122], 1X Glutamine [Life Technologies, #25030081], 0.1 mM 2-Mercaptoethanol [Life Technologies, #21985023], 1X B27 minus Vitamin A [Life Technologies, #12587010], 1X N2-Supplement-B [STEMCELL Technologies, #07156], and total cell number was established using manual cell counts of a 1:10 dilution in Trypan Blue Solution, 0.4% [Corning, #25-900-CI]. Cells were diluted in (DIV)0 medium (N2B27 supplemented with 10 µM Ascorbic acid (AA) [STEMCELL Technologies, #72132], 20µM SB431542 [Sigma, #S4217), 0.1µM LDN-193189 [Stemgent, #04-0074-02], 3µM CHIR 99021 [Tocris, #4423/10], 10 ng/mL FGF2 [R&D Systems, #233-FB], 10 µM RI [Tocris, #1254/50], and plated in ultra-low attachment plates [Corning, # 3471] at a concentration of 1 x 10^5^ cells/ml (for hESCs) or 2 x 10^5^ cells/mL (for iPSCs) in either 6-well (2ml/well) or 12-well (0.5ml/well) plates to promote the formation of Embryoid Bodies (EBs; visible on DIV1 under the microscope). Except when noted otherwise, for all protocols 100 nM Retinoic Acid (RA) [Sigma, #R2625] was added on DIV2, and 10-40 ng/ml BMP4 [R&D Systems, #314-BP-01M] was added on DIV4. Depending on the target sensory neuron subtype, 3 μM CHIR, 10µM DAPT [Tocris, #2634/10], 50 ng/ml Nerve Growth Factor (NGF) [Tocris, #256-GF], 20 ng/ml Brain Derived Neurotrophic Factor (BDNF) [Tocris, #212-BD/CF]; 10 ng/ml Glial Derived Neurotrophic Factor (GDNF) [Tocris, #212-GD/CF], or 25 ng/ml Neurotrophin 3 (NT3) [Tocris, #267-N3/CF], were added to the media on DIV2, DIV4, DIV7, DIV9, or DIV11 (as detailed in Supplemental Table S1). Up until DIV7, medium changes were performed by centrifuging EBs (4 min. at 200 RCF) and aspirating old media; after DIV7, EBs were transferred to 15 ml conical tubes, allowed to settle at the bottom, and spent media aspirated. On DIV 11 (“mechanoreceptor” and “proprioceptor” protocols) or DIV14 (“nociceptor” protocol), EBs were dissociated into single cell suspensions and plated as 2D cultures. To dissociate, EBs were collected in 15 ml conical tubes, washed once in PBS, and incubated in 1 ml 0.05% Trypsin [Gibco 25-300-054] supplemented with 25 µg/ml DNase I [Roche, #11-284-932-001] for 15 min at 37°C (shaking the tube every 5 minutes). EBs were mechanically dissociated by gentle trituration after which the digestion reaction was stopped with 2 volumes heat-inactivated fetal bovine serum (HI-FBS) [Life Technologies, #10082147] and 2 volumes of complete trituration wash media (CTWM) (PBS, 2.5% HI-FBS, 0.5 mM EDTA, 0.1% BSA [Life Technologies, #15260037], 25 mM Glucose [Sigma, #G8769], 2 mM MgCl2, 1X B27 minus Vitamin A, 1X N2 Supplement B). The single cell suspension was spun down for 5 min at 200 RCF, and resuspended in NDM sensory neuron media (Neurobasal [Life Technologies, #21103049], 1X P/S, 1X L-Glutamine, 1X NEAA and N2 Supplement B, 1X B27 minus VitA, 0.1 mM 2-Mercaptoethanol, 10 µM AA [Tocris, #4055] and 10ng/ml IGF-1 [R&D, #291-G1-200]). Cells were seeded on poly-ornithine (PO; 1:5 diluted in H2O to 20 μg/ml) [Sigma, #27378-490] and Laminin (1 mg/ml) [Life Technologies, #23017-015] or Cultrex coated plates at 50,000 cells/cm^2^. On the first day of plating, the media was also supplemented with 25 μM L-Glutamic Acid [Sigma, #G5889], as well as 1 μM Uridine (U) [Sigma, #U3750] and 1 μM Fluorodeoxyuridine (FDU) [Sigma, #F0503] to prevent proliferation of remaining undifferentiated cells. In addition, media was supplemented with the relevant neurotrophic factors depending on the target sensory subtype population being generated (see Supplemental Table S1 for details). NDM sensory neuron media was changed every 2-3 days. When using Laminin as culture substrate, NDM media was supplemented weekly with 1 μg/ml Laminin.

### Cloning of guide (g) RNA and donor plasmids

All gRNAs were designed using the Deskgen online algorithm (https://www.deskgen.com). Sequences of gRNAs used in this study are listed in Supplemental Table S2. Sense and antisense sequences were ordered as single-strand oligos, annealed, and cloned into the U6 gRNA expression vector (Addgene, #41824) (Mali et al., 2013) using either Gibson Assembly Mastermix (NEB, #E2611) or InFusion HD In-Fusion® HD Cloning Plus [Takara, #638920]. The *AAVS1* gRNA plasmid was obtained from Addgene [#41818] (Mali et al., 2013). *AVIL*, *NTRK1*, *MAFA*, and *RUNX3* gRNAs were tested using the T7 Endonuclease-I method as described previously (Garcia Diaz et al., 2020). Cas9 plasmids used in experiments were pCAGGS-Cas9-mCherry (Jacko et al., 2018), and pCAGGS-Cas9-mScarlet (Jacko, Wichterle, and Zhang; unpublished).

The *AVIL* donor plasmid was generated by amplifying a 1.98 kb fragment from RUES2 genomic DNA encompassing the *AVIL* stop codon and flanking areas (Supplemental Figure S1). Using this DNA template, *AVIL* left and right homology arms (*AVIL*-LHA and *AVIL*-RHA, respectively) were amplified with LHA and RHA specific primers (See Supplemental Table S3). The LHA-reverse primer contained a [G to A] substitution to mutate the PAM site in the donor plasmid. PCR reactions were performed using KAPA 2x HiFi PCR mix [KAPA Biosystems, #KK2602] or Promega 2x PCR mix [Promega, #M7505]; Gel purifications were performed with Zymoclean gel DNA purification kit [Zymo Research, #D4001]. Amplified and gel-purified PCR products of *AVIL*-LA, *AVIL*-RA, and P2A-iCre, were combined in the Afl-II predigested pCDNA3.1 donor plasmid by InFusion [Takara Bio, #638920]. The *AAVS1:loxSTOPlox:tdTomato* donor plasmid was described previously (Mali et al., 2013; Garcia Diaz et al., 2020).

The *NTRK1:tdT* donor was generated by amplifying left and right homology arms (*NTRK1*-LA, and *NTRK1*-RA respectively) from RUES2 hESC genomic DNA and cloned into pBSK-AscI by In-fusion. The *NTRK1*-LHA reverse primer included a BamH1 restriction site and an infusion homology area with the forward RHA primer. The forward RHA primers also contained the 3’ region of the tdTomato sequence that contains the BsrG1 site. The P2A-tdTomato reporter sequence was generated by cloning the BamH1-BsrG1 P2A-tdT fragment from a HB9:P2A-TdT reporter construct (Garcia Diaz et al., 2020) into the Ntrk1 donor plasmid digested with BsrG1 and partially with BamH1 (to avoid digesting the BamH1 site in the RHA).

*MAFA:tdT* and *RUNX3:tdT* donor plasmids were generated using a similar cloning strategy. For the *MAFA:tdT* donor, the homology arms (LHA and RHA) were amplified using InF-MAFA-F and InF-MAFA-R primers and cloned in the linearized pDonor plasmid by Infusion. The P2A:tdT fragment was amplified from *pDonor-AVIL:tdT* using InF-MafAtdT-F and InF-MafAtdT-R primers and cloned into the linearized pDonor-MAFA plasmid (using InF-pdMafA-F and InF-pdMafA-R primers) by Infusion. For the *RUNX3:tdT* donor plasmid, LHA and RHA were amplified from RUES2 hESC genomic DNA using the InF-Rx3-F and InF-Rx3-R primers and cloned into the linearized pDonor plasmid (using pdInverse-F and pdInverse-R primers) by InFusion. The pDonor-RUNX3 plasmid was linearized by PCR using InF-pdRx3-F and InF-pdRx3-R primers. The P2A:tdT fragment was amplified from a *pDonor-AVIL:tdT* plasmid (Oliver, Garcia Diaz and Corneo; unpublished result) using InF-Rx3tdT-F and InF-Rx3tdT-R primers and cloned to the linearized pDonor-RUNX3 through InFusion.

### Generation of hESC/iPSC reporter lines

hESC reporter lines were generated essentially as described previously (Garcia Diaz et al., 2020). In brief, hESCs were cultured to 70-80% confluency. On day of electroporation, media was refreshed with new media containing Rock Inhibitor, CloneR2, or CEPT. After at least one hour, cells were rinsed in PBS (w/o Mg^2+^ and Ca^2+^), incubated with Accutase [Sigma, #A6964] for 5 min at 37°C, dislodged from the plate by pipetting, and resuspended in 5 ml mTeSR™. Cell density was determined through manual cell counts, and for each electroporation, 2 x 10^6^ cells were spun down (5 min at 200 rcf) and resuspended in electroporation solution containing: 82 μl Nucleofection solution, 18 μl Supplement (both from the Amaxa P3 kit) [Lonza, #V4XP-3024), 7.5 μg gRNA plasmid, 2.5 μg Cas9-mCherry or Cas9-mScarlet plasmid, and 10 μg donor plasmid. Cell/DNA electroporation mixture was loaded into a cuvette and electroporated using a Nucleofector 4D electroporator using the hESC H9 program. Electroporated cells were incubated for another 15 min. in the electroporation solution and then diluted in 2 ml mTesR^TM^1 supplemented with CEPT or Rock Inhibitor and plated on Matrigel or Cultrex-coated plates in one well of a 6 well plate. After 24 hrs, the presence of Cas9 expression was assessed based on mCherry or mScarlet fluorescence. Cells were washed once with PBS (w/o Mg^2+^ and Ca^2+^), incubated with Accutase (5 min. at 37°C), dislodged from the culture plate and resuspended into a single cell suspension by pipetting. Cells were diluted with 5 ml of Fluorescent Activated Cell Sorter (FACS) media (10% KSR in PBS w/o Mg^2+^ and Ca^2+^) and centrifuged for 5 min at 200 rcf. Cell pellets were resuspended in 500 μl FACS media and passed through a 40 μm cell strainer. mCherry^+^ or mScarlet^+^ hESCs/iPSCs were isolated using either a Bio-Rad S3e sorter or a SONY MA900. The purified population was collected in mTeSR1 Plus with CEPT and seeded in Matrigel/Cultrex-coated plates at a density of 250-500 cells/cm^2^. Cells were maintained in mTeSR1 Plus and grown into distinct colonies for ∼10 days. Once formed, individual well-separated clones were picked by scraping of cells with a p200 micropipette tip and seeded into Matrigel- or Cultrex-coated U-bottom 96-well plates. When clones were sufficiently grown, each well was split (using 0.5 mM EDTA) into two 96 well plates, one used for genotyping and one for culture expansion. For genotyping, cells were washed with 200 μl PBS and incubated overnight in 100 μl Proteinase K (PK) lysis buffer (200 mM NaCl, 100 mM Tris-HCl pH 8.5, 5 mM EDTA, 0.2% SDS, supplemented with 0.1 mg/ml Proteinase K [PK; Sigma, #P2308]) in a humidified chamber at 56°C. The following day, PK was inactivated at 95°C and DNA was resuspended by pipetting and diluted 1:10 in H2O. Genotyping was performed by polymerase chain reaction (PCR) using 2x PCR mix [Promega, #M7505], 0.5 μl of (25 μM/μl) relevant primer mix (see Supplemental Table S4 for primer sequences), and 2 μl diluted clone DNA template in water (25 μl total reaction volume). Positive clones were identified based on the presence of Cre or tdT, and the appropriate band sizes for left arm (LA) and right arm (RA) insertion sites (see Supplemental Table S4, Figure 3, Supplemental Figures S1 and S5 for details). Positive clones were also assessed for heterozygous or homozygous reporter insertions. Clones with correctly inserted reporters were expanded and frozen down. Subsets of positive clones were further evaluated by differentiating the modified hESC/iPSC lines into sensory neurons and assessed for tdTomato expression.

### Immunohistological analyses

*EB staining.* On day of collection, EBs were washed once in PBS and fixed in 4% paraformaldehyde for 15 minutes on ice. Following fixation, EBs were washed in PBS, equilibrated in 30% sucrose for at least 2-3 hrs., and mounted in Tissue Tek OCT [O.C.T. Compound, FisherScientific, #4585] in Peel-A-Way molds [Polysciences, #18985-1]. EBs were sectioned at 30 μm on a cryostat [Leica CM1850]. Sections were stained in primary antibodies (see Supplemental Table S5 for details) in PBS, 1% BSA and 0.1% Triton X-100, overnight at 4°C. Following incubation in primary antibodies, sections were washed in PBS (2 x 6 minutes at RT) and incubated in secondary antibodies (see Supplemental Table S5) for 2 hrs. at RT. Sections were washed in PBS (2 x 6 minutes at RT) and cover-slipped with Vectashield [Vectorlab, #h-1000-10] or Fluromount [SouthernBiotech, #0100-01]. *Staining of 2D neuronal cultures.* Dissociated EB progenitor neurons were plated on coverslips (ThemoScientific, #3323) pretreated with 1 N nitric acid for 1hr, washed three times with water and sonicated for 1 h with 1 N HCl. After pretreatment, coverslips were washed 10 times with water and rinse twice with and maintained in 95% ethanol before use. Coverslips were coated with PO/Laminin, Matrigel, or Cultrex (all as described above) before plating cells. On day of collection, cells on coverslips were gently washed once in PBS and fixed in 4% Paraformaldehyde (in PBS) for 10 min at 4°C. Neurons on coverslips were stained with primary and secondary antibodies as described above. Images were acquired on LSM510 Meta, LSM700, or LSM900 (all Zeiss) confocal microscopes.

### Calcium imaging

Imaging was carried out using Fluo-4 calcium sensitive dye and epifluorescence microscopy as previously described (Bosco et al., 2024). Briefly, neurons were bulk-loaded with Fluo-4 AM, and imaged at 2 Hz for 60 s. After a 6 s control period, 30 mM KCl was locally applied to a field of cells for 2 s. Fluorescence intensity changes were measured from manually-drawn regions of interest encompassing neurons identified by their large round cell bodies or by tdTomato expression. ΔF/F was calculated by dividing the average pixel intensity of a ROI at each time-point by that of the baseline period prior to KCl addition.

### Electrophysiology

Action potential and passive membrane properties were assessed using conventional whole cell current clamp technique. DIV11 or DIV14 neurons were plated on 15 mm diameter coverslips at a density of 50,000 cells per well in a 24-well culture plate and maintained until ∼DIV28-32 prior to recording. Membrane potential recordings were performed using a Multiclamp 700B amplifier and a Digidata 1550 digital-to-analog converter. Signals were recorded at a 10-kHz sample rate using pClamp 10 software [all from Molecular Devices]. Patch pipettes were fabricated with a P-97 pipette puller [Sutter Instruments] using 1.5 mm outer diameter, 1.28 mm inner diameter filamented capillary glass [World Precision Instruments]. Pipette resistance was 2-5 MΩ when filled with the pipette solution. The external recording solution contained 145 mM NaCl, 5 mM KCl, 10 mM HEPES, 10 mM glucose, 2 mM CaCl2 and 2 mM MgCl2. The pH was adjusted to 7.3 using NaOH and the osmolality adjusted to 325 mOsm with sucrose. The pipette solution contained 130 mM CH3KO3S, 10 mM CH3NaO3S, 1 mM CaCl2, 10 mM EGTA, 10 mM HEPES, 5 mM MgATP and 0.5 mM Na2GTP (pH 7.3, 305 mOsm). Experiments were performed at room temperature (21–23°C). Correction for an empirically-calculated junction potential of –16 mV was applied prior to recording. During recordings, current was injected to hold the cells at −60 mV. Resting membrane potential was measured immediately following establishment of the whole-cell configuration. Membrane resistance and capacitance were calculated from the membrane potential changes in response to 1 s duration hyperpolarizing current steps that increased incrementally by 5 pA. The existence of H current was determined by the presence of a “sag” in the membrane potential greater than 10% evoked by a 1 s current injection which hyperpolarized the neuron to -80 mV. Single action potentials were evoked using a 2 ms duration current step. Trains of action potentials were evoked using 1 s duration depolarizing current steps that increased incrementally by 5 pA. An action potential was defined as a transient depolarization of the membrane which had a minimum rise rate > 10 mV/ms and reached a peak amplitude > 0 mV. Action potential characteristics were measured from the first action potential evoked from a 1 s current step at rheobase. The action potential duration was calculated from the full width at the half maximum voltage. For this calculation, the amplitude was measured from the threshold potential to the maximum potential. The maximum number of action potentials was measured from a 1 s current step. The amplitude of this step was dependent on the individual cell.

### Fluorescence activated cell sorting (FACS) of dissociated sensory neurons

DIV50 AVIL:tdT^+^ neuronal cultures were collected in ice-cold Hank’s balanced salt solution (HBSS) dissociated though enzymatic digestion using papain followed by collagenase/dispase (Malin et al., 2007; de Nooij et al., 2015). Papain solution consisted of 3 ml HBSS with ∼16 units/ml papain [Worthington, #LK003176], 0.83 mM L-cysteine, 0.42 mM EDTA, and 20 units DNAse I/ml [Roche, #4536282001]. Collagenase/dispase solution contained 3 ml HBSS containing 1,066 units/ml collagenase IV [Wortington, #LS004186], 4 units/ml dispase [Wortington, #LS02100], and 20 units/ml DNase I.

Incubation times were 10 min at 37°C for both enzymatic digestion steps. Solutions were exchanged by a low-speed centrifuge step (4 min. at 100 rcf) and aspiration. After collagenase/dispase digestion, enzyme solution was replaced with 500 μl HBSS supplemented with 20% HS and 20 units DNAse I, and DRG/cell suspension was dissociated by slow mechanical trituration using a p200 pipette. Cell suspension was passed through 40 μm gauze filters to clear remaining cellular aggregates. FACS of fluorescently-labeled (tdTomato^+^) neurons was performed at 12 psi, using a Becton Dickinson FACSAria, equipped with a 130 μm nozzle, and using a 586/15 (tdTomato) filter. Fluorescent neurons were plated in PDL/laminin-coated 96-well culture plates and imaged using a Plate RunnerHD system [Trophos].

### Semi-quantitative RT-PCR

RNA was isolated from hESCs/hiPSCs or differentiating EBs/neurons at various ages using the Qiagen RNAeasy miniprep kit [Qiagen, #74104] as per the manufacturer’s directions. In some cases, hESCs/hiPSCs, or EBs were stored in RNALater [Invitrogen, AM7020] prior to RNA isolation. First-strand cDNA was generated using the ThermoScientific RevertAid First Strand cDNA Synthesis Kit [Thermo Fisher Scientific, #K1691]. cDNA was diluted in water and used at 2 ng/semi-quantitative PCR reaction using 2x PCR mix [Promega, #M7505] and 0.5 μl of the relevant forward and reverse primer mix (each 10 μM; See Supplemental Table 6 for primer sequences). Reactions were allowed to proceed for 35 cycles and run on a 2% agarose gel before imaging.

### Neuronal counts and statistical analysis

Neuronal nuclei counts in EB tissue sections (30 μm) were performed using an ImageJ-based machine learning approach. This method follows four steps as summarized below.

*Image Acquisition and Preprocessing:* Confocal images of differentiating sensory neurons stained for nuclear markers (e.g. BRN3A, ISLET, SOX10, AP2α, DAPI) were captured using a Zeiss confocal microscope and data was exported as .czi or .lsm files, with each file consisting of a Z-stack of images. A maximum intensity projection (MIP) was applied to create a 2D TIFF composite file for analysis. *Channel Splitting:* Using ImageJ or Python, all confocal images were split into their respective red, green, and blue channels. Each channel was saved as an 8-bit grayscale image to enable further analysis. *Weka Segmentation for Classifier Creation:* Weka Trainable Segmentation was used to create classifiers for each channel/cells, with the blue channel serving as a classifier for DAPI (“all nuclei”), the red channel for one of the experimental nuclear markers (e.g. BRN3A or SOX10), and the green channel for the second experimental nuclear marker (e.g. ISL1 or AP2α). Yellow cells in the RGB image were used to classify co-expression of the experimental nuclear markers one and two (e.g. BRN3A and ISL1). The classifiers were trained on a sample dataset (>1000 cells) to segment cells accurately. *Segmentation and Cell Counting:* The trained classifiers were applied to their respective images to generate binary masks, separating positive cells from the background. Thresholding was used to convert the images into binary masks where cells were represented as white and the background as black. The Watershed algorithm was then employed to separate overlapping cells, ensuring they were segmented into distinct entities for accurate identification of positively stained cells. Cells were counted using ImageJ’s Analyze Particles function (equivalent to Python’s cv2.findContours), which identifies and counts individual particles (i.e. nuclei). To detect and categorize particles, Tukey’s method was used. The IQR, defined as the difference between the third quartile (Q3) and the first quartile (Q1), helped flag nuclei smaller than the lower bound (Q1 - 1.5 * IQR); these particles were removed, as they likely represented noise. The large (Q3 + 1.5 * IQR) particles were considered “fused” nuclei that failed to separate through the applied Watershed algorithm. To estimate the number of “fused” nuclei, the total surface area of the large particles was divided by the average nuclear size of the red and green nuclei. The total nuclear count consisted of the sum of the estimated single (unfused) nuclei and the estimated number of fused nuclei. For the dense DAPI staining, the total cell count was estimated by dividing the total area from the Analyze Particles plot by the average cell area of the red (e.g. BRN3A or SOX10) and green (e.g. ISL1 or AP2α) stained nuclei.

For counts of neuronal 2D cultures, all positive neurons (identified based on morphology or tdT expression) or positive nuclei (e.g. DAPI, BRN3A, ISLET, SOX10, or AP2α) in a given image were included in the count. Except when stated otherwise, averages were derived from a minimum of three biological replicates, each with 3-8 technical replicates. Graphs were generated using R. Average counts/values and standard error of the mean (s.e.m.) were calculated using Sigmaplot. Whenever used, Box plots in Figures show the median, 25th, and 75th percentile, and whiskers extend to the most extreme data point less than 1.5 times the interquartile range. Statistical analysis (Student’s t test or Mann– Whitney U test) on counted neuronal populations was performed using Sigmaplot. Significance was accepted for p<0.05 and denoted as * (p<0.05), or ** (p<0.01).

#### Resource availability

Requests for further information and resources should be directed to and will be fulfilled by the lead contact, Joriene de Nooij. All unique/stable reagents generated in this study are available from the lead contact with a completed materials transfer agreement.

## Acknowledgements

We thank Carmen Birchmeier (Max Delbruck Center, Berlin) for the cMaf antibody, and Marek Napierala (UT Southwestern), and the Friedreich’s Ataxia Cell Line Repository for the corrected FRDA4676 and FRDA68 lines. We also thank Luke Hammond for the discussion on the EB neuronal counting methodology, and Grace Shin and Callie Barber (both The Ohio State University) for critical reading of the manuscript. These studies were supported by the Thompson Family Foundation Initiative on CIPN, the Friedreich Ataxia Research Alliance (CU17-1234), and the CDMRP/DOD (HT94252310092) (all J.C. de Nooij).

## Author Contributions

EMG, KO, and SC performed sensory neuron differentiations and data analyses, EM, KO and AGD performed gene-editing of hPSCs, DW performed electrophysiological experiments, BC advised on the project, and JN conceived of the study, performed experiments and analyzed data. JN wrote the manuscript with input from all other authors.

