## Supplemental Materials for "Derivation and analysis of human somatic sensory neuron subtypes facilitated through fluorescent hPSC reporters"

**Supplemenal Information**

**Supplemental Table 1. Schematic of sensory neuron differentiation protocols.**

Growth and differentiation factors and small molecules generally used in the derivation of “nociceptive” (N), “low threshold mechanoreceptive” (M), or “proprioceptive” (P) sensory neuron subtypes. The final concentration, and the day of *in vitro* culture (DIV) on which each factor is added to the baseline media (N2B27 or NDM) are indicated.

**DIV0**

**DIV2**

**DIV4**

**DIV7**

**DIV9**

**DIV11**

**DIV14**

**DIV16-> ?**

N2B27

N2B27

N2B27

N2B27

N2B27

N2B27

NDM

NDM

AA

10μM

**N**

**M**

**P**

**N**

**M**

**P**

**N**

**M**

**P**

**N**

**M**

**P**

**N**

**M**

**P**

**N**

**M**

**P**

**N**

**M**

**P**

**N**

**M**

**P**

RI

10μM

**N**

**M**

**P**

FGF2

10ng/ml

**N**

**M**

**P**

SB

20μM

**N**

**M**

**P**

**N**

**M**

**P**

LDN

100nM

**N**

**M**

**P**

**N**

**M**

**P**

CHIR

3μM

**N**

**M**

**P**

**N**

**M**

**P**

**N**

**N**

DAPT

10μM

**M**

**P**

**M**

**P**

**N**

**M**

**P**

**N**

**M**

**P**

RA

100nM

**N**

**M**

**P**

BMP4

40ng/ml

**N**

**M**

**P**

NGF

50ng/ml

**N**

**N**

**N**

GDNF

10ng/ml

**N**

**M**

**N**

**M**

**N**

**M**

BDNF

20ng/ml

**M**

**M**

**M**

NT3

25ng/ml

**P**

**P**

**P**

B27-A

1X

**M**

**P**

**N**

**M**

**P**

**N**

**M**

**P**

IGF1

10ng/ml

**M**

**P**

**N**

**M**

**P**

**N**

**M**

**P**

bMOH

25μM

**M**

**P**

**N**

**M**

**P**

**N**

**M**

**P**

gluE

25μM

**M**

**P**

**N**

UFDU

1μM

**M**

**P**

**N**

**Small Molecules**

[final]

**Supplemental Table S2. Guide RNAs used in experiments.**

| **Locus** | **reporter** | **chromosome** | **gRNA sequence [5’-3’]^a^** | **# positive clones het/hom^b^** |
| --- | --- | --- | --- | --- |
| *AVIL* | iCre | 12 | TCCAAATGAAGAAAGAAAAG  GCTCCAAATGAAGAAAGAAA  GAAGGCCTATACCTATTGCA | hESC^c^: 22/2 |
| *AAVS1* | lxp-STOP-lxp:tdTomato | 19 | GGGCCACTAGGGACAGGAT | hESC: 12/0 |
| *NTRK1* | tdTomato | 1 | CTTGATGCTGTGGCGTTGC  CACGCCCGGCTGCAAGCCC  CCAGGACATCCAGGTAGAC | hESC: 17/4  hiPSC^d^: 4/7  FA-iPSC: 18/5 |
| *MAFA* | tdTomato | 8 | CACGGCCGACTTCTTCCTGT  CGGCGCCTACAGGAAGAAGT  CGACTTCTTCCTGTAGGCGC | hESC: 8/2  hiPSC: 5/7  FA-iPSC: 4/5 |
| *RUNX3* | tdTomato | 1 | GCGGGAGGAGTCCACCAGGG  GGCCCTACTGACCGCCCTGG  CCAGCGGGAGGAGTCCACCA | hESC: 10/4  hiPSC: 4/3  FA-iPSC: 1/1 |

^a^For each locus three guides were tested using T7E1 assays; gRNAs used in experiments are indicated in blue font. ^b^With exception of Avil:iCre, the number of colonies screened was 96; for Avil:iCre, 192 colonies were screened. ^c^hESCs used in experiments are RUES2. ^d^hiPSCs and FA-iPSCs correspond to FRDA patient (FA) and isogenic control iPSCs obtained from the Friedreich’s Ataxia Cell Line Repository at UT Southwestern (Li et al., 2016; Misiorek et al., 2020) and will be described elsewhere.

**Supplemental Table S3. Sequences of primers used for cloning donor plasmids.**

| **Target** | **primers** | **sequence [5’-3’]** |
| --- | --- | --- |
| AVIL LA and RA targeting area | Avilgenot-F-1  Avilgenot-R-3 | GGCACGGTGGCTCATACCTGTAATC  GCTTTGCGACACTGTGAACC |
| NTRK1 LA | Ntrk1-f2  LATA-P2A | GGTACTTTTTTGGCGCGCCGAGAAGAAATGGAATTAGTGC  gctcgtccatgccgtacagGGATCCGACAGGAGGTGCCTGGGCC |
| NTRK1 RA | Tomato-RATA  Ntrk1-r3 | ctgtacggcatggacgagctgtacaagtaaTACCTGGATGTCCTGGGC  TGGCGGCCTTTTTTGGCGCGCCAGCTAAGAAGAGTAACAGAGG |
| MAFA LA and RA targeting area | InF-MafA-F  InF-MafA-R | ACTCATCAATGTATCGCTACCAGCATCACCTCAACC  TTACAATTTACGCCTCGAGCGACTCATGCTGTGACT |
| RX3 LA and RA targeting area | InF-Rx3-F  InF-Rx3-R | ACTCATCAATGTATCAGGAAGAGGAGAGCCAGGTCTAG  TTACAATTTACGCCTGGAAAGCCAACAGTTAGGAAC |

**Supplemental Table S4. Primers used for genotyping gene-edited clones.**

| **donor** | **segment** | **primers** | **sequence [5’-3’]** |
| --- | --- | --- | --- |
| *AVIL-P2A-Cre* | AVIL - LA | AVIL-gtF1  P2A-R1 | GGCACGGTGGCTCATACCTGTAATC  GCTTTAACAGAGAGAAGTTCGTGGC (1059 bp) |
|  | AVIL - RA | P2A-F1  AVIL-gtR3 | GCCACGAACTTCTCTCTGTTAAAG  GCTTTGCGACACTGTGAACC (2063 bp) |
|  | AVIL-genot (wt/het/hom) | AVIL-F1  AVIL-R1 | AACAAGCAAGGAATGCAAGG  GAACTCCCTCTTGCTCCAACTT (wt: 373 bp; Cre^+^: 1498 bp) |
| *NTRK1-P2A-tdT* | NTRK1-genot (LA) | NTRK1-F1  P2Agenot-R4 | CCTCCTAATGCAGTCTGCTCC  CATAGGTCCAGGGTTCTCCTCC (1029 bp) |
|  | NTRK1-genot (RA) | TAgt-F1  TAgt24-R3 | AGAGCAGAAATGAATCCTGG  ATGGGACAGGGAGATGGCTT (wt: 1190 bp; tdT+: 2670 bp) |
|  | NTRK1-genot (wt/het/hom) | TAgt-F1  TAgt-R1 | AGAGCAGAAATGAATCCTGG  ATAATTGCTATGACGGGACC (wt: 552 bp; tdT^+^2046 bp) |
| *MAFA-P2A-tdT* | MAFA-genot (RA) | MAFtdT-F  MAFA-R3 | GCAGACTTCTTCCTGGGATCCGGAGCCACG  GGTTGAGCATAGTAGGACAGC (2332 bp) |
|  | MAFA  (wt/het/hom) | MAFtdT-F  MAF-T7E1-R1 | GCAGACTTCTTCCTGGGATCCGGAGCCACG  CCTGGTGTCCACGTCCTGTA (wt:123 bp; tdT^+^: 1613bp) |
| *RUNX3-P2A-tdT* | RX3-genot (LA) | RX3-2F  P2Agenot-R1 | GCAGGTCTGTGTTCGGAGTC  GCTTTAACAGAGAGAAGTTCGTGGC (762 bp) |
|  | RX3-genot (RA) | tdT-F1  RX3-2R | ATGGTGAGCAAGGGCGAGGAGG  CCTGCCAAGAGAACAGAGAGT (2225 bp) |
|  | RX3- genot (wt/het/hom) | RX3-2F  RX3-2R | GCAGGTCTGTGTTCGGAGTC  CCTGCCAAGAGAACAGAGAGT (wt 1525 bp; tdT^+^ 3018 bp) |
| tdT | tdT-genotyping | tdT F1  tdT R3 | ATGGTGAGCAAGGGCGAGGAGG  GCTGAAGGGCGAGATCCACCA (490 bp) |
| P2Aseq-1 | NTRK1 |  | GGAAGCGGAGCTACTAACTTCAGCCTGCTGAAGCAGGCTGGCGACGTGGAGGAGAACCCTGGACCT |
| P2Aseq-2 | AVIL, MAFA, RX3 |  | GGATCCGGAGCCACGAACTTCTCTCTGTTAAAGCAAGCAGGAGACGTGGAAGAAAACCCCGGTCCT |

**Supplemental Table S5. Antibodies and Dyes used in Experiments.**

| **Antibody** | **Dilution** | **Source and cat#** | **RRID** |
| --- | --- | --- | --- |
| Rb anti-hAdvillin | 1:100 | Abcam #ab175126 | AB_2861264 |
| Rb anti-mAdvillin | 1:500 | Abcam #ab72210 | AB_1951510 |
| M anti-Ap2a | 1:50 | DSHB #3B5 | AB_528084 |
| M anti-Brn3a | 1:200 | Millipore #mab1585 | AB_94166 |
| Gp anti-Islet1/2 | 1:20,000 | Tanabe et al., 1989 |  |
| M anti-islet 1/2 | 1:100 | DSHB #39.4D5 | AB_2314683 |
| GP anti-cMAF(N term.) | 1:5000 | C. Birchmeier (#2223) |  |
| M anti-Pax3 | 1:50 | DSHB #Pax3 | AB_528426 |
| Rb anti-Peripherin | 1:800 | Millipore #AB1530 | AB_10681273 |
| M anti-Runx3 | 1:200 | Santa Cruz #101553 | AB_2184397 |
| Rb anti-Sox2 | 1:500 | ThermoFisher #481400 |  |
| Gt anti-hSox10 | 1:400 | R&D Systems #AF2864 | AB_442208 |
| Gp anti-dsRed | 1:30,000 | T.M. Jessell CU1906 |  |
| Rb anti-RFP | 1:1000 | Rockland #600-401-379-RTU |  |
| Rb anti-mTrkA | 1:5000 | L. Reichardt |  |
| M anti-TrkB | 1:200 | Santa Cruz #377217 |  |
| M anti-Tuj1 | 1:1000 | R&D Systems #MAB1195 | AB_357520 |
| **Dyes** | | |  |
| DAPI | 1:1000 | ThermoFisher #D1306 |  |
| NeuroTrace | 1:1000 | Mol. Probes #N-21483 |  |

.

**Supplemental Table S6. Sequences of primers used for RT-PCR.**

| **Target** | **primers** | **sequence [5’-3’]** |
| --- | --- | --- |
| NGN1 | hNGN1-f  hNGN1-r | TCCCCCTCCCCTAGTCAGCA  GGCGACCTAACAAGCGGCTC |
| NGN2 | hNGN2-f  hNGN2-r | GGCCAAAGTCACAGCAACGCT  CGATCCGAGCAGCACTAACACG |
| HOXC5 | hHOXC5-f  hHOXC5-r | CCCGCCACAGATTTACCCGT  CAGAGTCTGGTAGCGCGTGT |
| HOXC8 | hHOXC8-f  hHOXC8-r | AGTAGCGAAGGACAAGGCCACT  CGGCTGTAAGTTTGCCGTCC |

**Supplemental Figure Legends**

**Supplemental Figure S1. A CRISPR/Cas9 gene-editing strategy to label human pluripotent stem cell derived somatic sensory neurons.**

A) Expression of Advillin and Islet (Isl) in p0 mouse DRG.

B) Expression of ADVILLIN and ISLET in adult human DRG.

C) Transcriptionally-defined human DRG neuronal subtypes and expression levels of *ADVILLIN* across all subtypes. Data obtained from the human DRG transcriptomic atlas (Nguyen et al., 2021).

D) Genomic location, guide RNA sequence (blue font), and PAM site (black bold font and underlined) to insert a P2A-Cre driver into the 3’UTR of the *AVIL* gene. STOP codon is marked in red font. Primer locations to genotype successful transgenic clones are indicated. For details on methods, guide RNA, or primer sequences, see Methods and Supplemental Tables S2-4.

E) Genotyping results for a correctly targeted *AVIL* locus. PCR results indicate correct targeting of the left arm (left panel), right arm (middle panel), and the presence of a homozygous Cre coding sequence insertion (right panel).

Scale: 20 μm (A), and 50 μm (B).

**Supplemental Figure S2. BMP4 promotes dorsal PAX3^+^ progenitor fates and increases sensory neuron yields.**

A, B) Schematic diagrams describing the main growth factors/cytokines and their time points of application to coach human ES cells towards a somatic sensory neuron fate. In b) Bone morphogenetic protein (BMP4) is added at DIV4 to promote dorsalization of neural progenitors.

C, D) Expression of SOX2 and PAX3 in DIV9 EBs cultured without (c) and with (d) addition of BMP4 at DIV4. Progenitor cells are co-labeled by the general neural marker Neurotrace (NT). Scale: 20 μm.

E, F) Expression of BRN3A and ISLET in DIV14 EBs cultured without (e) and with (d) addition of BMP4 at DIV4. Progenitor cells are co-labeled by the general nuclear marker DAPI. Boxes in images on the left indicate area enlarged shown in images on the right. Scale: 20 μm.

G) Percentage of ISLET^+^, BRN3A^+^, and BRN3A^+^ISLET^+^ sensory neurons at DIV14 when cultured without (gray boxes) or with (orange boxes) the addition of BMP4 at DIV4. Data derived from four independent differentiations with at least three representative EB images analyzed per data point (differentiation). Median percentage (± s.e.m.) of ISLET^+^ neurons is 36.42 (± 9.79) in absence of BMP4, and 43.18 (± 5.83) in the presence of BMP4 (p=0.424, two tailed t-test). Median percentage of BRN3A^+^ neurons is 6.57 (± 3.49) in absence of BMP4, and 44.93 (± 9.38) in the presence of BMP4 (p=0.009, two-tailed t-test) Median percentage of BRN3A^+^, ISLET^+^ neurons is 3.72 (± 2.27) in absence of BMP4, and 19.47 (± 7.06) in the presence of BMP4 (p=0.038, two-tailed t-test). Black open circles represent data obtained from independent differentiations with CHIR added from DIV0-7, BMP4 (10 ng/ml) from DIV2-4, DAPT from DIV9-11, and all GFs (NGF, GDNF, BDNF, NT3; 10 ng/ml) starting on DIV9 or 11 and onwards. Blue circles represent data obtained from independent experiments from the same starting ESCs (with/without BMP4). Red circles represent data from a differentiation with 5 ng BMP4 added on DIV4 (instead of 10 ng).

H) Expression of *HOXC5*, *HOXC8*, *NGN1*, and *NGN2* in developing EBs between DIV0 and DIV14 as determined by semi-quantitative RT-PCR. Similar results were obtained from two independent experiments.

**Supplemental Figure S3.** **Differentiation efficiencies across different human ESC and iPSC lines.**

A) Expression of BRN3A and ISLET in DIV14 EBs from the *H9* human ESC and *FA0000011* iPSC lines. For *H9*, EBs are co-labeled by Neurotrace (in blue). Boxes in images on the top row indicate area enlarged shown in images on bottom row. Scale 20 μm.

B) Percentage of BRN3A^+^,ISLET^+^ sensory neurons in DIV14 EBs across different human ESC and iPSC lines. Mean percentage (± s.e.m.) for *H9* hESC is 23.2% (data derived from one experiment with four representative EB images analyzed. Mean percentage for *Fa0000011* is 7.9% (± 0.2%) (data derived from two independent experiments with at least four representative EB images analyzed per data point). Mean percentage for FA68 is 40.8% (± 4.2%) (data derived from five independent experiments with at least three representative EB images analyzed per data point). Mean percentage for FA4676 is 26.6% (± 8.4%) (data derived from four independent experiments with at least three representative EB images analyzed per data point).

**Supplemental Figure S4.** **Electrophysiological analyses of hESC-derived sensory neurons.**

A) Membrane resistance of tdT^+^ and tdT^off^ *AVIL:tdT* hESC-derived sensory neurons. Black open circles represent neurons derived from EBs treated with 10 or 40 ng/ml BMP4 at DIV4-7. Neurons derived from EBs treated with 40 ng/ml BMP4 at DIV4-7 are further segregated in tdT^off^ (open red circles) and tdT^+^ (solid red circles) sensory neurons. Solid black circles represent neurons derived from a differentiation protocol with a shortened CHIR (DIV0-4), the inclusion of RA (100nM at DIV2-4), 40 ng/ml of BMP4 (DIV4-7), and earlier DAPT treatment (DIV4-14).

B) Action potential (AP) trace recorded from a hESC-derived sensory neuron. The action potential is evoked by a 2 ms current injection, represented by the lower trace. The AP trace is overlaid on a subthreshold trace in which the current injected was insufficient to trigger an AP. An arrow indicates a shoulder on (i.e. a slowing of) the falling phase of the action potential.

C) Percentage of neurons with an H-current or a distinguishable shoulder (S) on the downward stroke of the action potential. Plot shows the average for all neurons per differentiation protocol and type: BMP 10 ng/ml (black open circle), BMP 40 ng/ml (gray closed circle), BMP 40 ng/ml tdT^off^ (open red circles), BMP 40 ng/ml tdT^+^, and RA (closed black circle).

D, E) K^+^-induced calcium transients in hESC-derived sensory neurons. Images of neurons under control conditions (left panel) and following addition of potassium chloride (30 mM KCL; right panel) are shown in (d). In (e), graph showing the average increase in Fluo-4 fluorescence intensity across time following the addition of KCL (marked by green bar). Scale in d: 20 µmb) Example trace of a hESC-derived sensory neuron with a shoulder on the downward slope of the AP.

F) FACS profile of dissociated AVIL:tdT^+^ sensory neurons.

G) FACS-isolated AVIL:tdT^+^ sensory neurons can be replated for morphological axon out-growth studies.

**Supplemental Figure S5. Characterization of a *MafA:tdTomato* and *Runx3:tdTomato* human iPSC reporter lines**

A, B) Genomic location, guide RNA sequence (blue font), and PAM site (black bold font and underlined) to insert a P2A-tdT reporter into the 3’UTR of the *MAFA* (A) or *RUNX3* (B) locus. Primer locations to genotype transgenic clones, and genotyping results are indicated. STOP codon is indicated in red font. Note in (B) that original STOP codon is mutated (gray font) and a new one generated by mutating the PAM site. For details on methods, guide RNA, or primer sequences, see Methods and Supplemental Tables S2-4.

C) Expression of the neural crest markers AP2α and SOX10 in DIV11 EBs.

D) Expression of the post-mitotic sensory neuron markers BRN3A and ISLET in DIV11 EBs.

E, F) Percentage of BRN3A^+^ (B), ISLET^+^ (I), and BRN3A^+^ISLET^+^ (B/I) neurons from the total number of cells in DIV11 EBs. Percentages are estimates based on a machine learning approach. See Methods for details. Data derived from 3 (DIV11) independent differentiations with at least 4 representative EB images used per data point (differentiation). Open black circles in (E) represent data from differentiations with CHIR on DIV0-4, RA (100 nM) on DIV2-4, BMP4 (40 ng/ml) on DIV4-7, DAPT on DIV4-14, and all GFs (NT3 at 25 ng/ml; BDNF at 20 ng/ml; GDNF at 10 ng/ml; NGF at 50 ng/ml) from DIV11 and onwards. Closed black circle in (E) is similar to open black circles but with NT3, BDNF, and GDNF (each 10 ng/ml; no NGF) starting on DIV7 and onwards. Open black circles in (F) represent data from differentiations similar to open black circles in (E) but with GFs (NT3 at 25 ng/ml; BDNF at 20 ng/ml; no GDNF or NGF) starting at DIV7 instead of DIV11.

G) Expression of tdT and RUNX3 in a cluster of sensory neurons migrating from a DIV16 EB exposed to differentiation media conditioned with skeletal muscle differentiation media.

H) Expression of tdT in DIV44 and DIV51 sensory neurons derived from *RUNX3:tdT* and *MAFA:tdT* hESCs respectively. After DIV30, hESC-derived neurons often cluster to form organoid-like or DRG-like structures.

Scale: 20 μm (C, D, G), 200 μm (H).

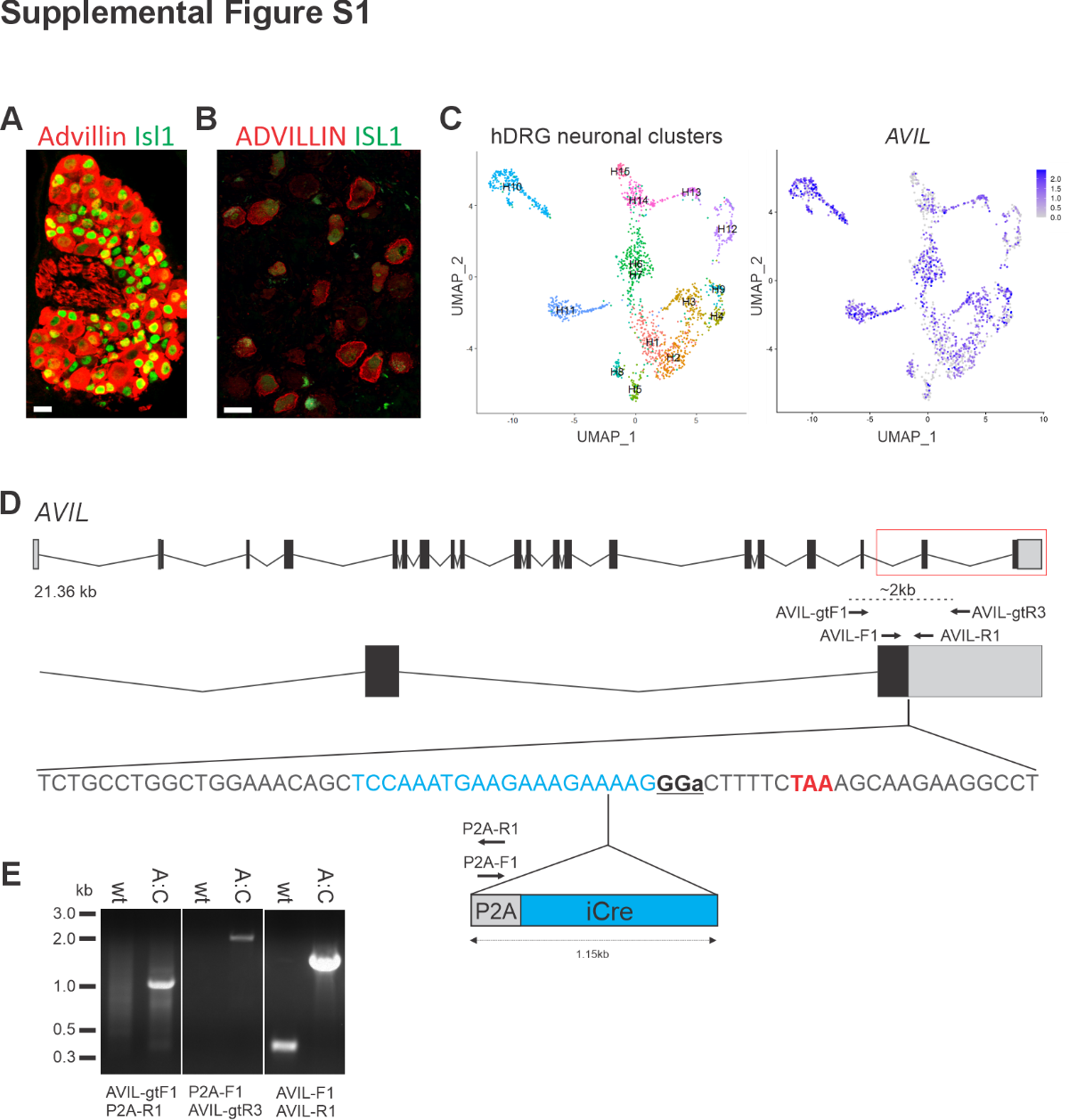

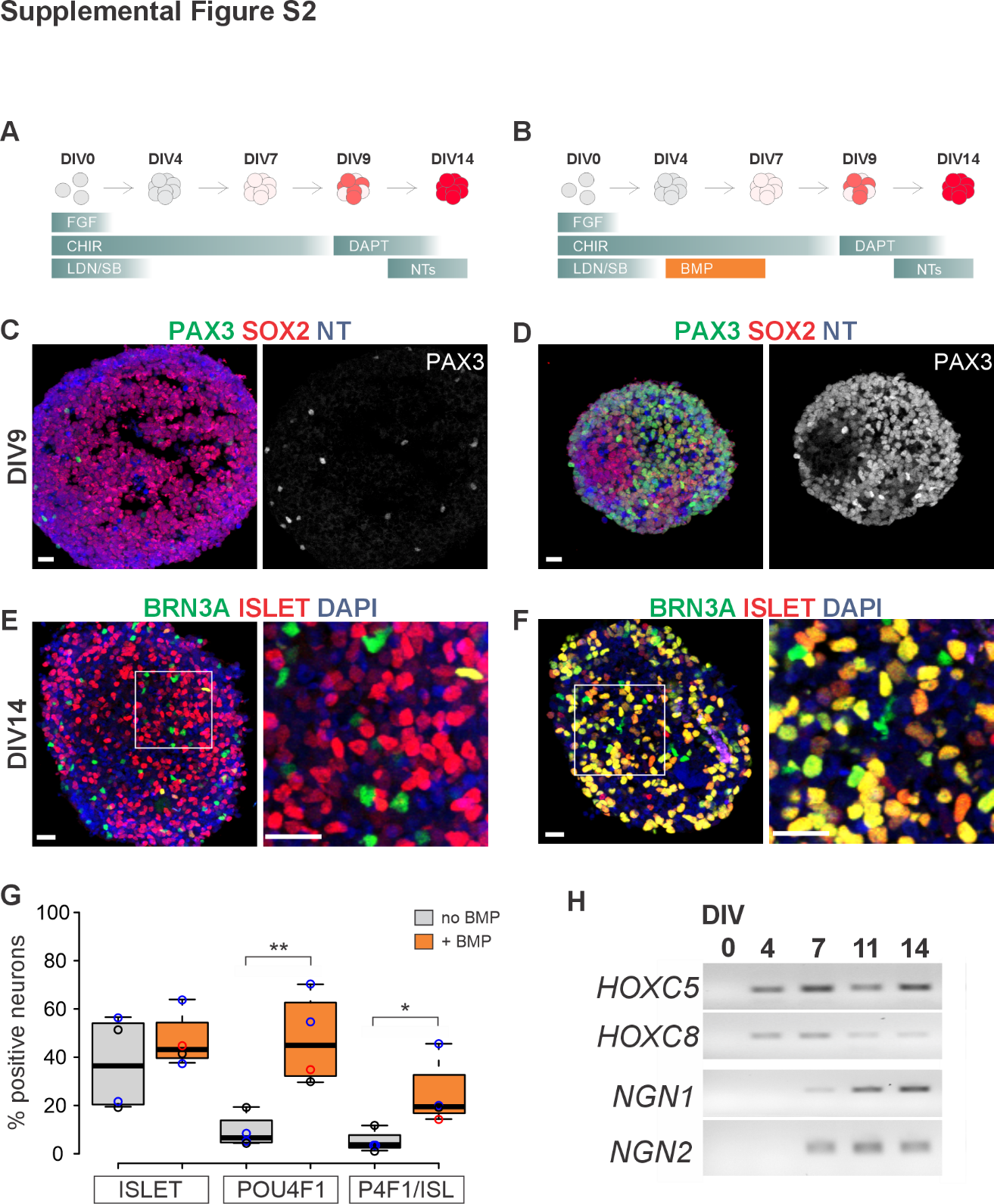

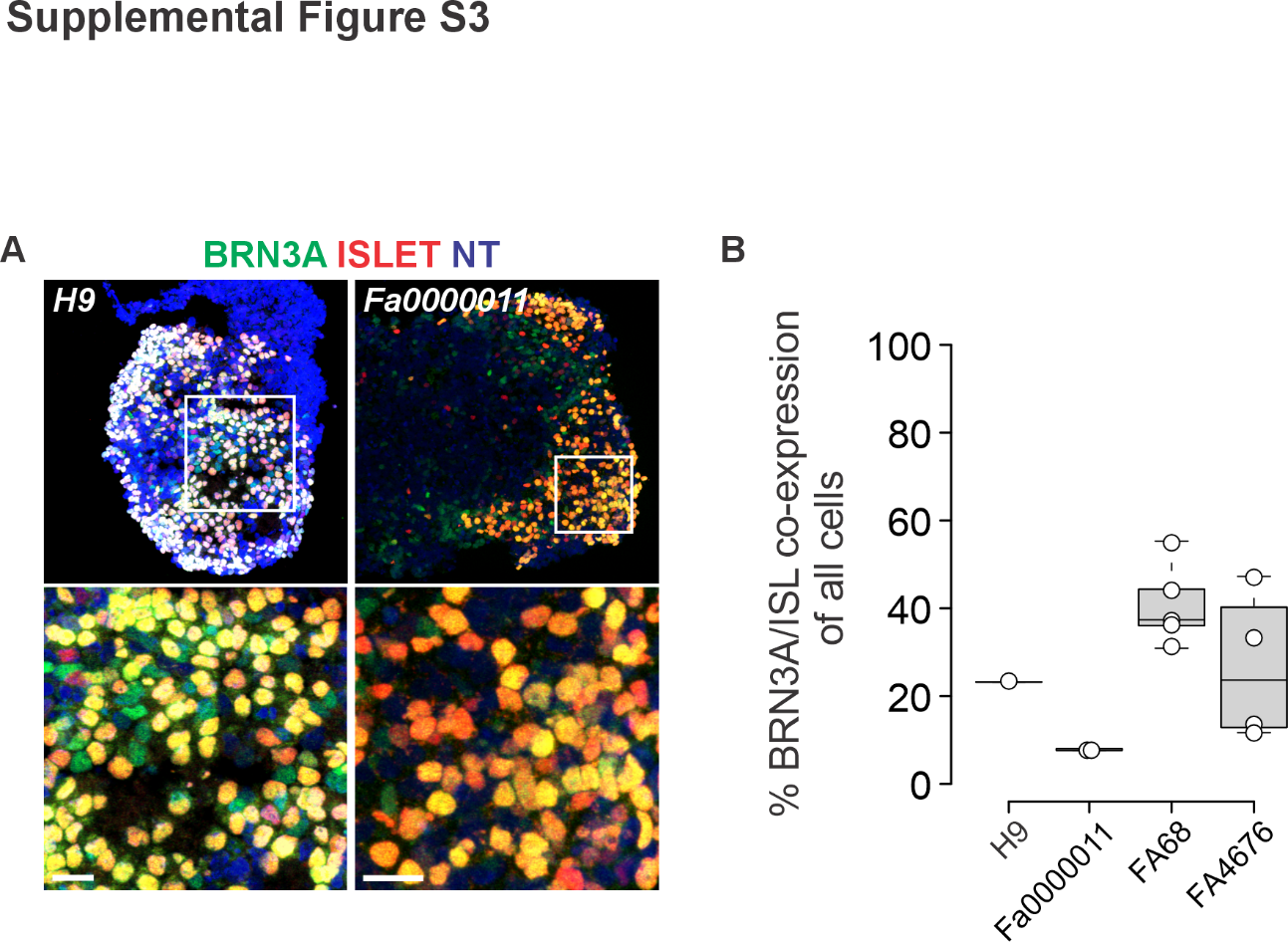

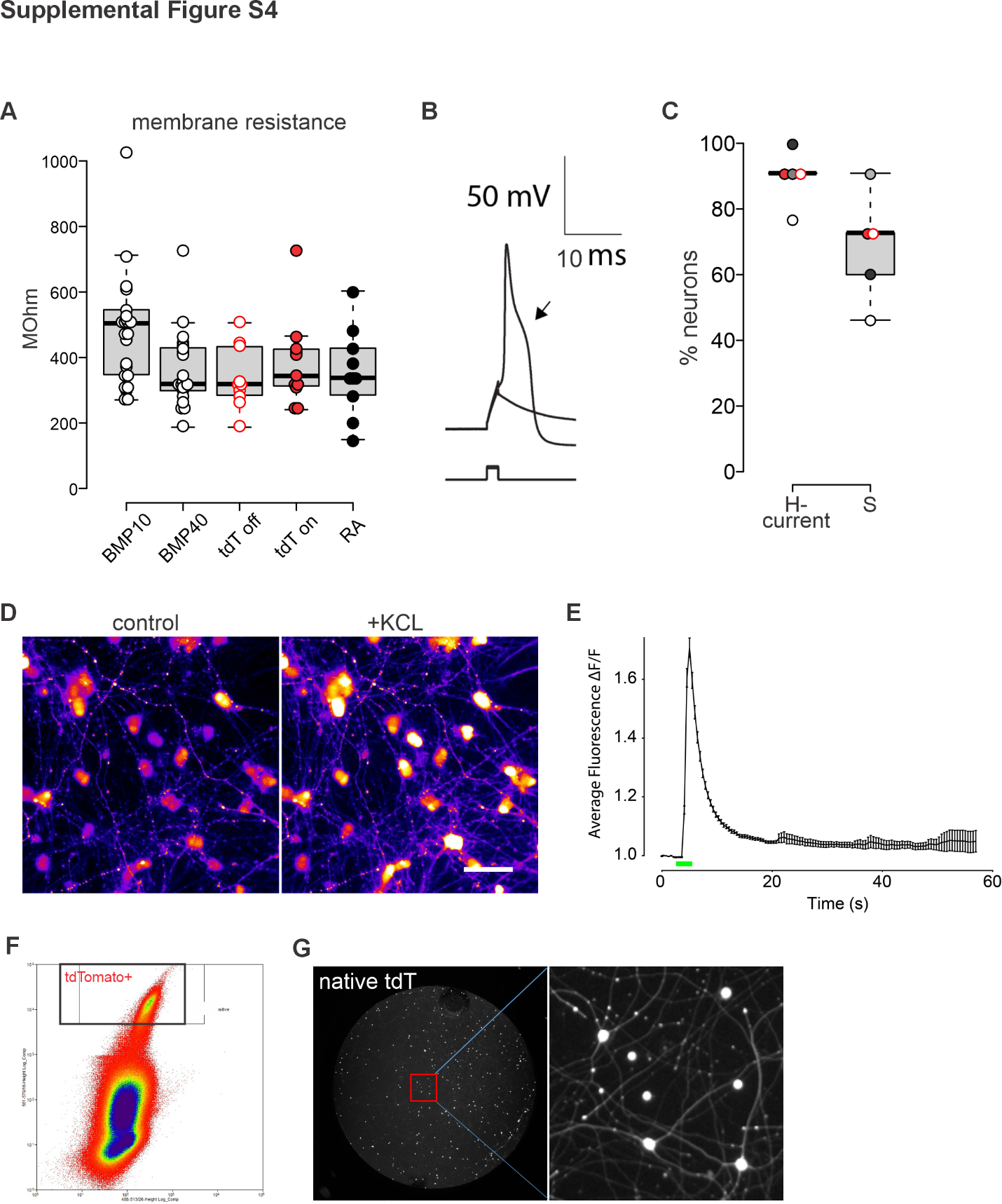

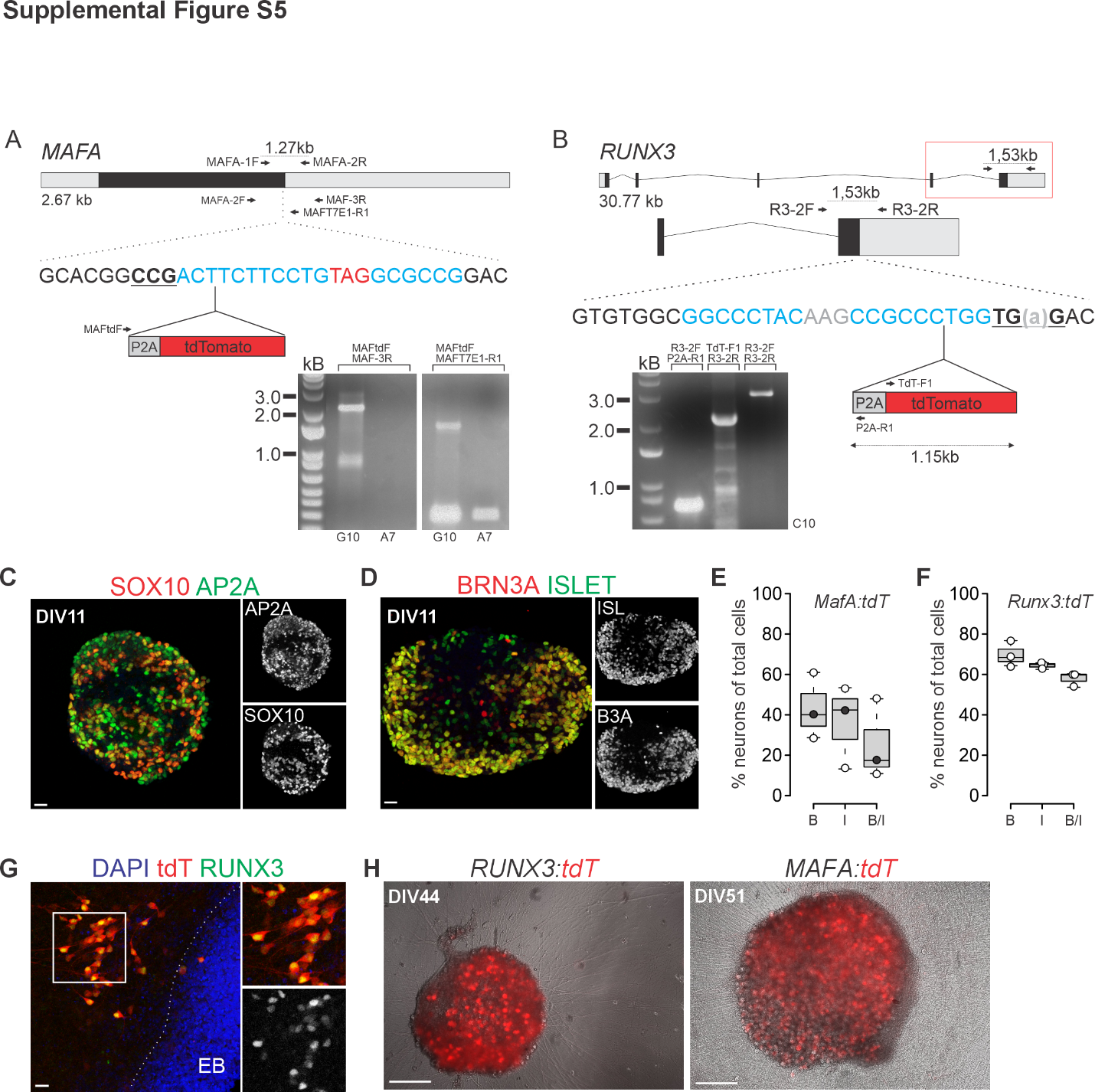
